# Emergence of Task-Related Motor Cortical Dysfunction in Mice with Progressive Parkinsonism

**DOI:** 10.64898/2026.05.29.728524

**Authors:** Hiba Douja Chehade, Daniil Berezhnoi, Samhitha Somavarapu, Zihang Wang, Jitong Jiang, Liqiang Chen, Benjamin B. Risk, Thomas Wichmann, Hong-Yuan Chu

## Abstract

The primary motor cortex (M1) exhibits substantial changes in Parkinson’s disease (PD). However, the temporal relationship between midbrain dopamine (DA) neurodegeneration, M1 circuit dysfunction, and motor symptoms remains poorly understood. Using the MitoPark mice, we investigated the time course of M1 cellular dysfunction during progressive DA loss. MitoPark mice developed gradually impaired skilled movements at a moderate motor stage of Parkinsonism. The motor deficits were associated with reduced cellular activity, impaired neuronal coactivation, and decreased proportion of responsive M1 pyramidal neurons. Moreover, the corticospinal (CSp), but not the intratelencephalic (IT), neurons showed a selective reduction in cellular activity during reaches. At the population level, M1 pyramidal neurons included heterogeneous functional clusters in relation to skilled movements and their stability in longitudinal encoding of movement degraded substantially in MitoPark mice. Together, these results delineate the emergence of M1 circuit pathophysiology at cellular and neural population levels during progressive Parkinsonism.

## INTRODUCTION

Parkinson’s disease (PD) is a chronically progressive neurological disorder associated with a reduced number and amplitude of spontaneous movements. These hypokinetic motor signs are causally linked with the degeneration of dopamine (DA) neurons in the substantia nigra pars compacta (SNc) and the accompanying DA depletion throughout the basal ganglia (*1, 2*). According to the traditional model of the pathophysiology of PD, the loss of DA neuromodulation has opposing effects on the activity of direct and indirect pathways of the basal ganglia, resulting in abnormally increased basal ganglia GABAergic inhibition of their downstream targets, including the thalamocortical network and the brainstem (*1–4*). In addition, the loss of DA neuromodulation induces complex anatomical and functional adaptations at the cellular and synaptic levels in the striatum (*5–7*), the globus pallidus (*8, 9*), and the subthalamic nucleus (*8, 10–12*). Overall, these changes promote the emergence of aberrantly synchronous burst patterns of neural activity throughout the basal ganglia and the thalamocortical network (*1, 2, 6, 13*). Compelling evidence suggests that the suppression of abnormally synchronized network activity is correlated with the therapeutic effects of deep-brain electrical stimulation (*2, 14*), supporting its pathological role in the pathophysiology of PD.

Patients with PD not only exhibit overall slowness of movement (i.e., bradykinesia) but also deficits in fine motor skills (e.g., micrographia) (*15*). Interestingly, in PD patients, impairment in manual dexterity likely occurs independently of other clinical measures of disease severity (*16*), indicating that the mechanisms underlying impairment in motor skills and general motor activity may not entirely overlap. It is known that the acquisition and execution of fine motor skills are highly dependent on cerebral cortical function (*16, 17*). Relevant to the pathophysiology of PD, the primary motor cortex (M1) and the supplementary motor area and premotor cortices are major sources of synaptic input to the basal ganglia, and they are also the primary downstream targets of basal ganglia GABAergic inhibition via the thalamus. In the parkinsonian state, M1 exhibits anatomical and functional alterations (*18–25*). Functional imaging studies have reported altered cortical activity and abnormal cortical plasticity in PD patients (*25, 26*). Animal model studies have provided additional mechanistic insight into cortical dysfunction following DA loss, including reduced firing rate of subsets of cortical neurons (*18, 22*), enhanced synchronous firing (*21–23, 27*), and abnormal movement encoding (*23, 28*).

Most prior preclinical studies on cortical circuit dysfunction were conducted using subacute or chronic neurotoxin models of fully developed Parkinsonism. These models do not adequately recapitulate the chronic, progressive nature of PD or exhibit robust intersubject variability in disease staging, making it impossible to capture stage-dependent changes in cortical circuits. Therefore, it is necessary to define the temporal relationship between dopaminergic degeneration, changes in cortical activity, and abnormalities in skilled movements in a chronically progressive model of the disease to better understand cortical dysfunction in PD.

In the present work, we used a progressive mouse model of nigrostriatal degeneration (i.e., MitoPark mice (*29–31*)) to conduct longitudinal cellular imaging of neural activity in the M1 of freely moving animals during gradually worsening striatal DA depletion. Our results demonstrate that M1 neurons gradually started to show abnormal activity at both cellular and population levels during the progression of Parkinsonism, with the corticospinal (CSp) neurons showing more robust task-related disruptions than the intratelencephalic (IT) neurons. These studies also showed that the magnitude of movement-related activity changes in M1 pyramidal neurons was greater in MitoPark mice than that in controls. Furthermore, we demonstrated that, as a whole population, M1 pyramidal neurons comprised distinct functional clusters for movement encoding, whose organization is disrupted as Parkinsonism develops in MitoPark mice. Together, the present work provides new insight into M1 circuit pathophysiology during the progression of parkinsonism.

## RESULTS

We used MitoPark mice as a model of progressive nigrostriatal dopaminergic neurodegeneration for longitudinal investigation of M1 circuit dysfunction and its relationship to parkinsonian motor deficits (*30*). We trained MitoPark mice and littermates to acquire and execute a self-initiated single-pellet reaching-and-grasping (SPRG) task (Fig. 1A, Key Resource Table 1) (*32*), which is recognized as a cortex-dependent skilled movement in rodents (*17, 33, 34*). Following the successful acquisition of the task, mice were subjected to *in vivo* GCaMP6f imaging to monitor M1 pyramidal neurons during the SPRG task (20 min/session/week, Fig. 1A) using a miniature endoscope (Inscopix). Given the well-characterized time course of neurodegeneration and behavioral changes in MitoPark mice (*29, 30, 35*), we selected 14-, 16-, 18-, and 20-week-old animals to define the temporal relationship between striatal DA depletion and cortical activity changes and its potential correlation with the development of skilled movement deficits at different stages of Parkinsonism.

**Figure 1.**
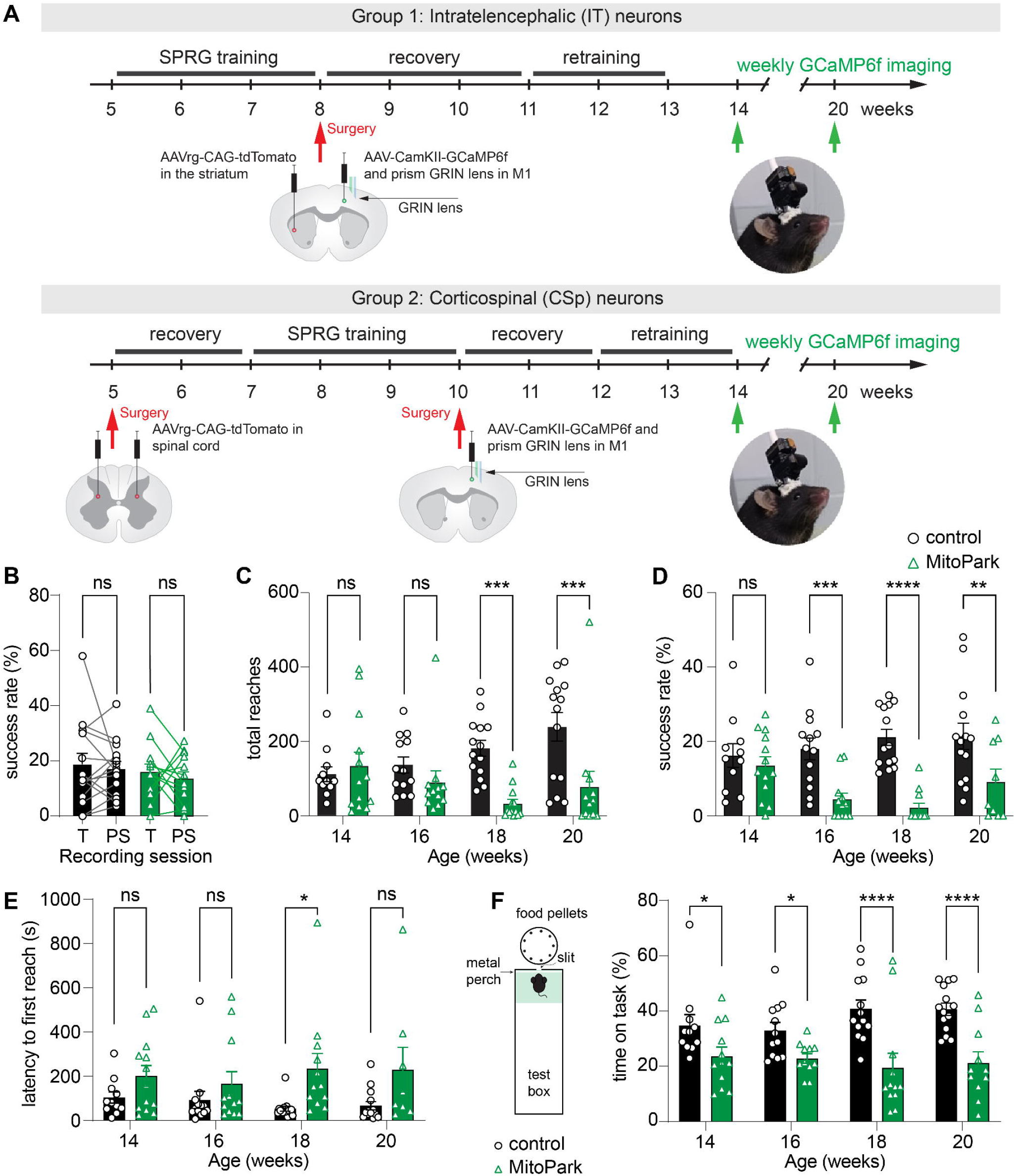
Skilled movement deteriorates during progressive loss of midbrain DA neurons. **A)** Diagram showing the overall experimental design. **B-D)** Performance of MitoPark mice and controls in the SPRG task, as measured by the success rate of control and MitoPark mice between the last day of training (T) and the first post-surgery (PS) recording session (B). Wilcoxon paired test. Total number of reaches (C) and success rate (D) significantly decrease in MitoPark mice compared to littermate controls. **E-F**) The latency to the first reach (E) and the time on task (F) were quantified to assess the task engagement for MitoPark mice and littermate controls. Linear mixed models were used to compare MitoPark versus control across weeks (see Methods for details). Each data point represents one mouse. MitoPark = 13, 12, 13, 12 mice and control = 11, 14, 14, 14 mice at 14, 16, 18 and 20 weeks, respectively. Results are reported as mean ± SEM hereafter. Significance codes: * p < 0.05, ** p < 0.01, *** p < 0.001, **** p < 0.0001. Detailed numbers and statistics are available in the Source Data Table for all figures.

**Table 1.** Key Resource Table.

### Skilled movement deteriorates during progressive loss of midbrain DA neurons

We trained MitoPark mice and littermate controls to learn the SPRG task beginning at 5 or 7 weeks of age (depending on the experimental groups, Fig. 1A). Both MitoPark mice and littermate controls could learn the SPRG task, performing different categories of reach-to-grasp movements (Fig. S1A). While the success rate of animals from both groups seems relatively low compared to prior work (Fig. 1B, D) (*36–40*), such discrepancies are likely due to technical details, including animals’ age at training (*41*). All trained mice, regardless of their success rates, from both groups were subjected to stereotaxic surgery, receiving gradient refractive index (GRIN) lens implantation in M1 at 8-10 weeks of age. Once recovered from surgery, animals were retrained for 2 weeks and assess for their ability to perform the skilled movements (Fig. 1A). We found that there was no difference in the success rates before and after the implantation surgery for both controls and MitoPark mice (Fig. 1B). These data suggest that the capacity to execute the skilled movements was preserved following the surgery.

Next, we studied animals’ performance in the SPRG task during progressive Parkinsonism. Relative to littermate controls, MitoPark mice showed comparable numbers of reaches and success rates at 14 weeks of age; however, the success rates started to decline as early as 16 weeks of age (Fig. 1C, D). Although these changes indicate that MitoPark mice developed impaired motor skills (e.g., the decreased success rate), it may also reflect changes in motivation and task engagement due to striatal DA depletion (e.g., the changes in the number of reaches).

To assess task engagement, we quantified the latency to initiate the first reach. While overall reach latency did not change systematically over time in either group, control mice showed a trend of shorter latency to start the first reach than MitoPark mice and this difference was statistically significant at 18 weeks of age (Fig. 1E). Moreover, we quantified the time on task (Material & Methods) as a measure of motivation and engagement (Fig. 1F). While there was no systematic change across time in either group, MitoPark mice spent less time on task relative to littermate controls across ages (Fig. 1F). These data suggest that, although both controls and MitoPark mice maintained their motivation and engagement in the task across sessions, MitoPark mice likely developed decreased motivation and impaired task engagement as early as 14 weeks of age. A comparison of the progressive changes in animals’ performance (Fig. 1C, D) and motivational impairments (Fig. 1E, F) indicates that the decreased success rates of MitoPark mice mainly reflect impaired motor skills, partially involving moderate motivational deficits.

To further assess impaired motor skills in MitoPark mice, we analyzed different categories of failed reaches. We found an increased percentage of missed reaches in MitoPark mice relative to controls, becoming statistically significant at 16 weeks of age and beyond (Fig. S1D). However, changes in proportions of drop (i.e., the mouse successfully grasped the pellet, but lost it during the retrieval phase) and flick (i.e., the mouse touched the pellet but failed to grasp it) reaches were not changed across time between controls and MitoPark mice (Fig. S1B, C). We further tracked forepaws of a subset of animals at 16 weeks of age with moderate Parkinsonism (N = 3 per group) and quantified task-related movement kinematics (Fig. S2A-C). We found that both MitoPark mice and littermate controls performed stereotyped and rapid reach-to-grasp movements (i.e., mean duration = 400 ms), as measured by strong across-trial correlations for both successful and missed reaches (Fig. S2D, E). It indicates that the task had been consolidated in the motor control circuits. We next calculated the reach kinematic for comparison between control and MitoPark mice. For successful reaches, MitoPark mice showed a notable reduction in reach trajectories (28% shorter) and the speed of reaches (∼20% slower, Fig. S2F-G) relative to controls. For missed reaches, there was no noticeable difference in forepaw trajectory length or movement speed between MitoPark mice and littermate controls (Fig. S2H-I). Together, these results indicate that altered kinematics may contribute to the impaired skilled movements in Parkinsonism.

### Decreased cortical neuronal activity during progressive Parkinsonism

M1 circuit is important for the execution of complex skilled movements (*33, 34, 42, 43*) and its network remodeling is linked with the impaired motor function in mouse models of Parkinsonism (*28, 40*). Thus, we next studied how the activity changes of the cortical neurons contribute to the impaired motor skills in MitoPark mice. To do this, we combined SPRG with *in vivo* miniscope imaging (Fig. 2A) for a longitudinal analysis of M1 neuronal activities in MitoPark mice and littermate controls. To this end, we injected AAV9-CaMKII-GCaMP6f into the deep layers of the caudal forelimb region of M1, followed by a subsequent implantation of a GRIN lens at the same location of AAV injections (Fig. 2A, B). Layer 5 (L5) pyramidal neurons connect the cortex with the striatum and subcortical motor regions, contributing to the execution of movement (*44–48*). Post-mortem histological analyses showed that (1) GCaMP6f expressed robustly in the forelimb region of M1 (*49*) (Fig. S3A, B); (2) the implantation consistently centered at the L5 of M1, corresponding to location of the retrogradely labelled CSp neurons via spinal injection of AAVrg-tdTomato (Fig. S3A-C), (3) there were no differences in the rostrocaudal, mediolateral, or dorsolateral coordinates of the bottom edges of the GRIN lenses in control and MitoPark mice (Fig. S3D-F); and (4) MitoPark mice showed a substantial reduction in the striatal tyrosine-hydroxylase (TH) positive axon terminals at the end of recording (21-24 weeks of age), confirming striatal dopamine loss (Fig. S3G-I). GCaMP6f signals were recorded and extracted from M1 pyramidal neurons in 14-, 16-, 18-, and 20-week-old MitoPark mice and littermates. We then quantified the rate and amplitude of Ca^2+^ events as indirect measures of neuronal activity and compared them across behavioral states between MitoPark mice and littermate controls. First, we identified “non-reaching epochs” in the recorded videos, defined as periods lasting at least 30 sec during which animals did not perform reaches. Remaining epochs during which animals performed active reaches were then defined as active “reaching epochs” (Fig. 2D). The duration of reach epochs declined gradually as SNc DA neurons progressively degenerated in MitoPark mice, compared with littermate controls (Fig. 2E), consistent with the reduction of the total number of reaches from early to late stages of MitoPark mice (Fig. 1C).

**Figure 2.**
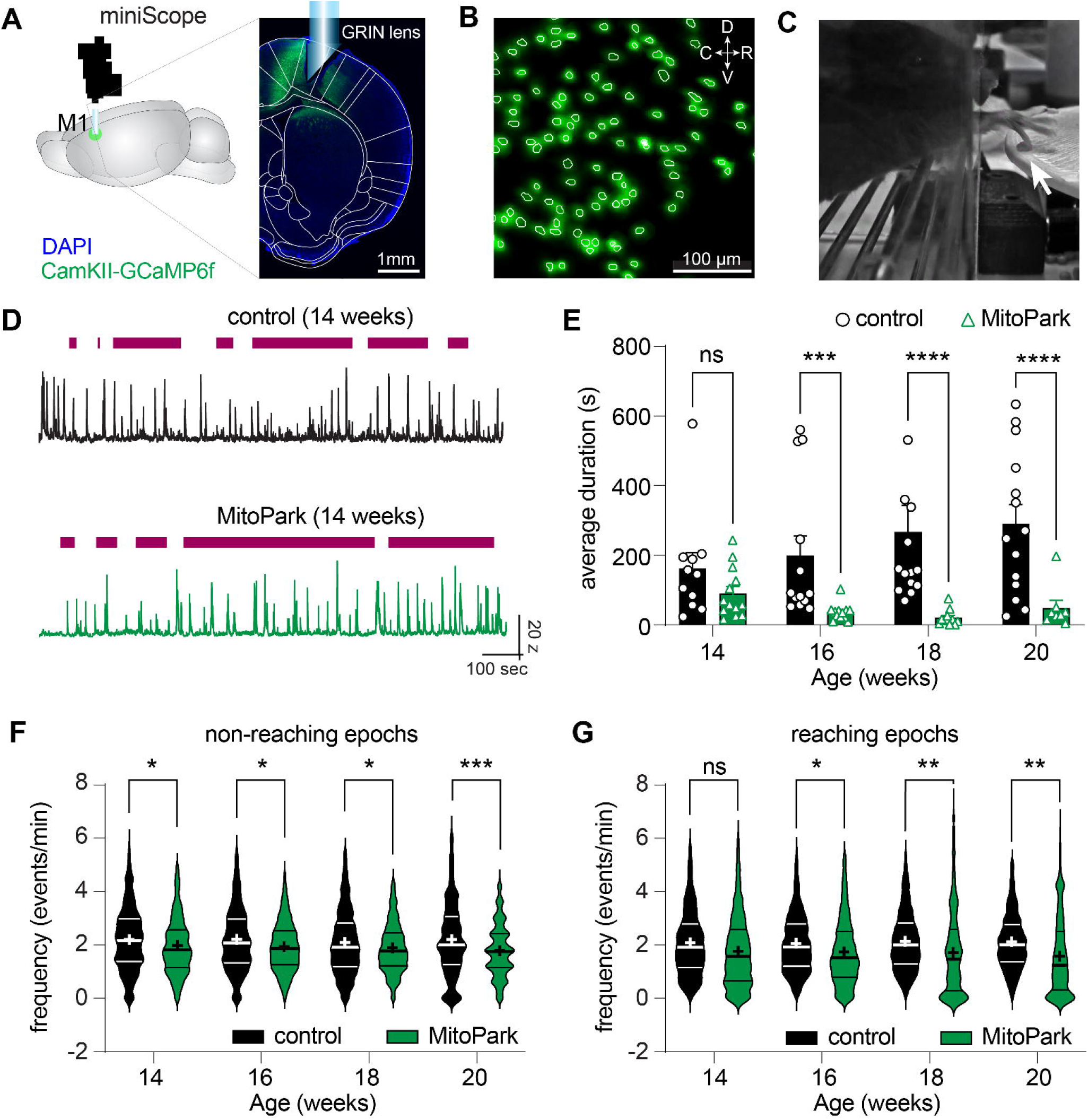
Decreased cortical pyramidal neuronal activity during progressive Parkinsonism. **A)** Diagram showing in vivo Ca2+ imaging from M1 and a representative histological image showing GCaMP6f expression and lens placement in M1. **B)** Maximal projection image of Ca2+ transients from a segmented field of view of the Prism GRIN lens. **C)** An example behavior video frame while a mouse was performing reach-to-grasp task, highlighting the grasp onset (white arrow). **D)** Extracted GCaMP6f traces of a representative neuron from control (top) and MitoPark (bottom) mice at 14 weeks. Horizontal lines on top indicate the reaching epochs. **E)** Duration of reaching epochs in MitoPark mice and littermates across ages. Linear mixed-effects model on log-transformed duration (see Methods for details). Each data point represents one mouse. Control = 11, 13, 14 and 13 mice, MitoPark = 13, 12, 12 and 8 mice, at 14, 16, 18 and 20 weeks, respectively. **F-G)** Frequency of Ca2+ events from M1 pyramidal neurons during non-reaching (F) and reaching (G) epochs in MitoPark mice and littermate controls. Horizontal lines indicate median and interquartile range (IQR), and cross (+) symbols indicate the mean. Linear mixed effects models were fit to compare behavioral states and within each behavioral state (see Methods for details). Violin plots represent cells. (F) Control n/N = 1203 cells/11 mice, 1385 cells/13 mice, 1699 cells/14 mice and 1610 cells/14 mice, MitoPark n/N = 1074 cells/13 mice, 1459 cells/12 mice, 2072 cells/11 mice and 1677 cells/12 mice, (G) Control n/N = 1191 cells/11 mice, 1354 cells/13 mice, 1712 cells/14 mice and 1612 cells/14 mice, MitoPark n/N = 1052 cells/13 mice, 1431 cells/12 mice, 1298 cells/11 mice and 1268 cells/9 mice at 14, 16, 18 and 20 weeks, respectively.

During non-reach epochs, animals were in behavioral states presumably less dependent on the motor cortex (*46, 50*). We found a significant group effect in the frequency of Ca^2+^ events (p = 0.0003). Specifically, the frequency of spontaneous Ca^2+^ events decreased by 12% at 14 weeks, 11% at 16 weeks, 10% at 18 weeks, and 20% at 20 weeks in MitoPark mice compared with age-matched littermate controls (Fig. 2F). Similarly, there was a main group effect during the reach epochs (p = 0.0009), which became statistically significant at 16 weeks and beyond (Fig. 2G). In the linear-mixed effect regression analysis, we detected a significant behavioral state x group interaction (p =0.027). Thus, although M1 pyramidal neuronal activity decreased in both reaching and non-reaching epochs, the extent of reduction was greater in reaching epochs than that in non-reaching epochs. We also quantified the amplitude and area under the curve (AUC) of individual Ca^2+^ events as additional measures of neuronal activities. During either non-reach or reach epochs, we found no difference in either the amplitude (Fig. S4A-C) or the AUC (Fig. S5A-C) of Ca^2+^ events between MitoPark mice and littermate controls. Together, these data demonstrate that the activity of M1 pyramidal neurons, as a population, decreased gradually and significantly from the early to the advanced motor stages in Parkinsonism, and that these changes were more pronounced when skilled movements were conducted.

### Cell-subtype-specific decreases in neuronal activity in Parkinsonism

Considering the cellular heterogeneity of cortical pyramidal neurons, we next examined whether subtypes of M1 pyramidal neurons are differentially affected by loss of DA, as reported previously (*18, 22, 23*). Specifically, we focused on the CSp and IT neurons in layer 5 that connect M1 with the basal ganglia circuits (Fig. 3A, E). CSp and IT neurons were retrogradely labeled by injection AAVrg-CAG-tdTomato into the spinal cord and contralateral striatum, respectively. *In vivo* dual-color imaging of GCaMP6f and tdTomato signals allowed us to monitor tdTomato-labeled CSp or IT neurons (Fig. 3B, F) by quantifying their activity changes during the development of Parkinsonism.

**Figure 3.**
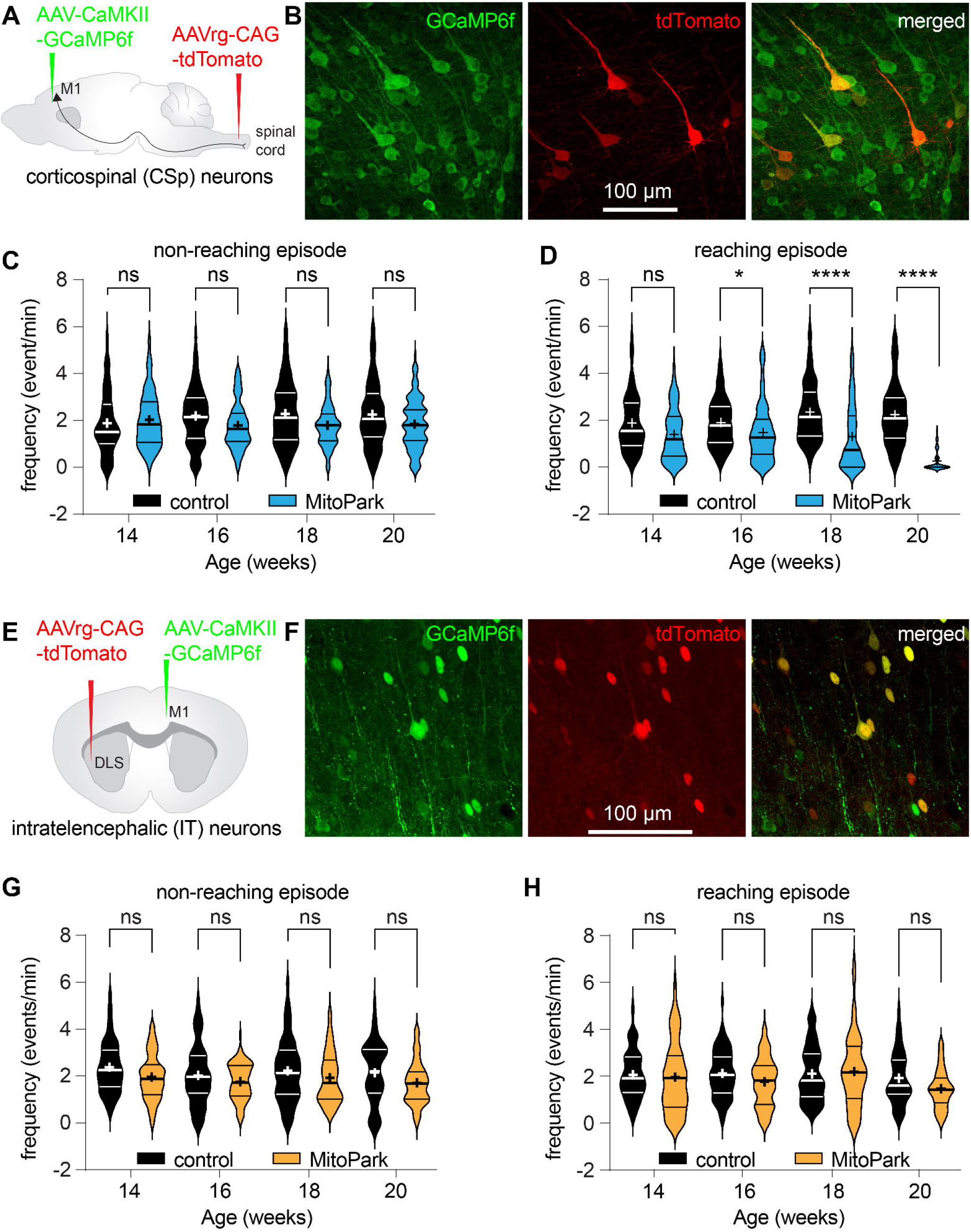
Cell-subtype-specific decreases in neuronal activity in Parkinsonism. **A)** Strategy to express GCaMP6f in all pyramidal neurons in M1 and retrogradely label CSp neurons. **B)** Representative images showing AAV expression in M1. Middle panel shows CSp neurons labeled with tdTomato in M1 ipsilateral to the injection site in the spinal cord. **C-D)** Frequency of Ca2+ events during non-reaching (C) and reaching epochs (D) from M1 CSp neurons. **E)** Strategy to express GCaMP6f in all pyramidal neurons in M1 and retrogradely label IT neurons. **F)** Representative images showing AAV expression in M1. Middle panel shows IT neurons labeled with tdTomato in M1 contralateral to the injection site in the DLS. **G-H)** Frequency of Ca2+ events from M1 IT neurons during non-reaching (G) and reaching (H) epochs. Horizontal lines indicate median and IQR, cross (+) symbols indicate the mean. Violin plots represent cells. A linear mixed effect model was fit for each behavioral state (see Methods for details). (C) Control n/N = 83 cells/5 mice, 189 cells/6 mice, 242 cells/6 mice and 177 cells/6 mice, MitoPark n/N = 138 cells/7 mice, 175 cells/7 mice, 171 cells/7 mice and 191 cells/7 mice, (D) Control n/N = 76 cells/5 mice, 179 cells/6 mice, 240 cells/6 mice and 177 cells/6 mice, MitoPark n/N = 129 cells/7 mice, 166 cells/7 mice, 134 cells/6 mice and 87 cells/4 mice, (G) Control n/N = 78 cells/6 mice, 63 cells/7 mice, 131 cells/7 mice and 50 cells/7 mice, MitoPark n/N = 88 cells/6 mice, 81 cells/5 mice, 108 cells/6 mice and 58 cells/5 mice, (H) Control n/N = 78 cells/6 mice, 63 cells/7 mice, 132 cells/8 mice and 49 cells/8 mice, MitoPark n/N = 89 cells/6 mice, 82 cells/5 mice, 51 cells/3 mice and 58 cells/5 mice, at 14, 16, 18 and 20 weeks, respectively.

We analyzed Ca^2+^ events of both neuronal subtypes during the non-reach and reach epochs separately. In non-reach epochs, the frequency of Ca^2+^ events in CSp or IT neurons was comparable between control and MitoPark mice across ages (Fig. 3C, G). In contrast, while the animals were actively reaching, CSp neurons exhibited an age-dependent reduction in Ca^2+^ event frequency in MitoPark mice relative to littermate controls (Fig. 3D), but no such alteration was observed in the IT neurons between groups (Fig. 3H). Specifically, CSp neurons showed comparable frequencies of Ca^2+^ events between 14-week-old MitoPark mice and littermate controls. CSp neurons from MitoPark mice showed a progressive decline in the frequency of Ca^2+^ events, with reductions of 22%, 45%, and 88% at 16-, 18-, and 20-weeks of age, respectively, relative to littermate controls (Fig. 3D). In both reach and non-reach epochs, there were no significant changes in the amplitude (Fig. S4D-G) or the AUC (Fig. S5D-G) of Ca^2+^ events in either CSp or IT neurons from MitoPark mice relative to littermate controls.

We conclude that M1 exhibited a cell-type-specific reduction in pyramidal neuronal activity in MitoPark mice as Parkinsonism developed. The hypoactivity of M1 CSp neurons, but not IT neurons, is partially consistent with predictions from the traditional model and prior electrophysiological studies in non-human primates (NHPs) (*22, 23*).

### Disrupted functional connectivity between M1 pyramidal neurons in Parkinsonism

Spatially distributed neuronal ensembles emerge in M1 during the acquisition of complex skilled movements (*33, 34, 42, 43*) and dynamic changes in M1 ensembles have been linked with the impaired motor function in a mouse model of Parkinsonism (*40*). Thus, we assessed coordinated activities of M1 pyramidal neurons from MitoPark mice and littermate controls across ages. By Z-scoring the observed co-activated Ca^2+^ events to that obtained from circularly shifted data (see Materials & Methods), we quantified the proportion of co-activated pyramidal neurons (regardless of PT or IT neurons) across different behavioral states in both groups (Fig. 4). Linear mixed effect model showed group main effects for both non-reaching and reaching epochs in the proportion of pyramidal cell pairs with Z ≥ 2 (the vertical line in Fig. 4A, B) between MitoPark mice and controls (Fig. 4C-D). This result indicates a moderate reduction in the percentage of co-activated cortical pyramidal neuron pairs in MitoPark mice relative to controls. However, the current study was not properly powered to determine subtype-selective changes in co-activated cell pairs within CSp and IT neurons. Together, we found that MitoPark mice appear to have fewer simultaneously activated pyramidal neurons in M1 relative to controls. Thus, the abnormally synchronized spontaneous electrical activity of M1 pyramidal neurons seen in Parkinsonism was absent in the present results (*21*) (likely due to the slow kinetics of GCaMP6f signals, Fig. S5A).

**Figure 4.**
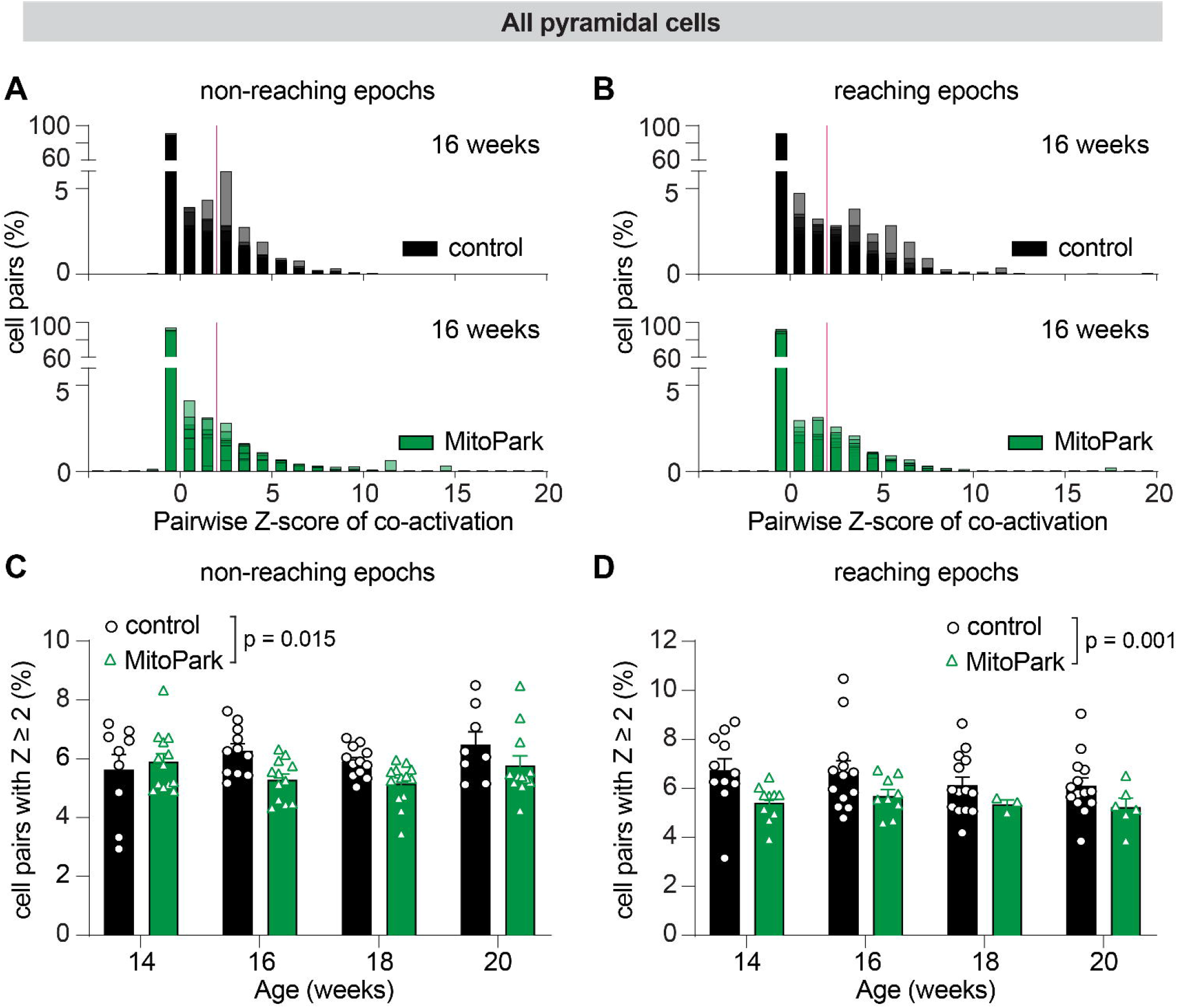
Disrupted functional connectivity between M1 pyramidal neurons in Parkinsonism. A-B) Distribution of pairwise Z-score of coactivation in control (top) and MitoPark (bottom), during non-reaching (A) and reaching (B) epochs. The vertical red line represents the threshold (Z ≥ 2) chosen to identify highly coactivated cell pairs. **C-D)** Proportion of co-activated cell pairs for nonspecific pyramidal neurons during non-reaching (C) and reaching (D) epochs. Linear mixed-effect model (see Methods for details). Group by age interaction was non-significant so the model was reduced to main effects of age and group, which showed significant group main effect for both non-reaching (p = 0.015) and reaching epochs (p = 0.001). (C) Control = 9, 11, 12, 8 mice and MitoPark = 13, 12, 13, 12 mice, (D) Control = 11, 13, 14, 14 mice and MitoPark = 11, 9, 3, 6 at 14, 16, 18 mice and 20 weeks, respectively.

### Impaired movement encoding by M1 pyramidal neurons in Parkinsonism

To further characterize the relationship between cortical activity changes and impaired skilled movements at cellular levels in progressive Parkinsonism, we performed peri-movement analyses by aligning *z*-scored Ca^2+^ signals (see Materials & Methods) to the onset of the grasps, which could be reliably identified from the behavior videos (Fig. 2C). We started this analysis with all GCaMP6f-expressing M1 pyramidal neurons (regardless of whether they were CSp or IT neurons). For the peri-movement analysis, we statistically compared GCaMP6f signals within a time window of ±1.0 s at the onset of grasps. Based on the direction of GCaMP6f changes, M1 neurons were classified into up- or down-modulated groups (Fig. 5A, B) corresponding to those with statistically increased or decreased amplitude of Ca^2+^ signals within the ±1 sec time window. M1 neurons that showed unaltered magnitudes of Ca^2+^ signals at the onset of grasp were classified as the non-modulated group (Fig. 5C) (*45*). 56%-64% of M1 pyramidal neurons in control mice across ages were characterized as non-modulated. This proportion exceeded 75% in 14- and 16-week-old MitoPark mice and > 80% in the 18- and 20-week-old groups (Fig. 5D). These data demonstrate that the proportion of movement-modulated cortical pyramidal neurons decreases gradually during progressive Parkinsonism.

**Figure 5.**
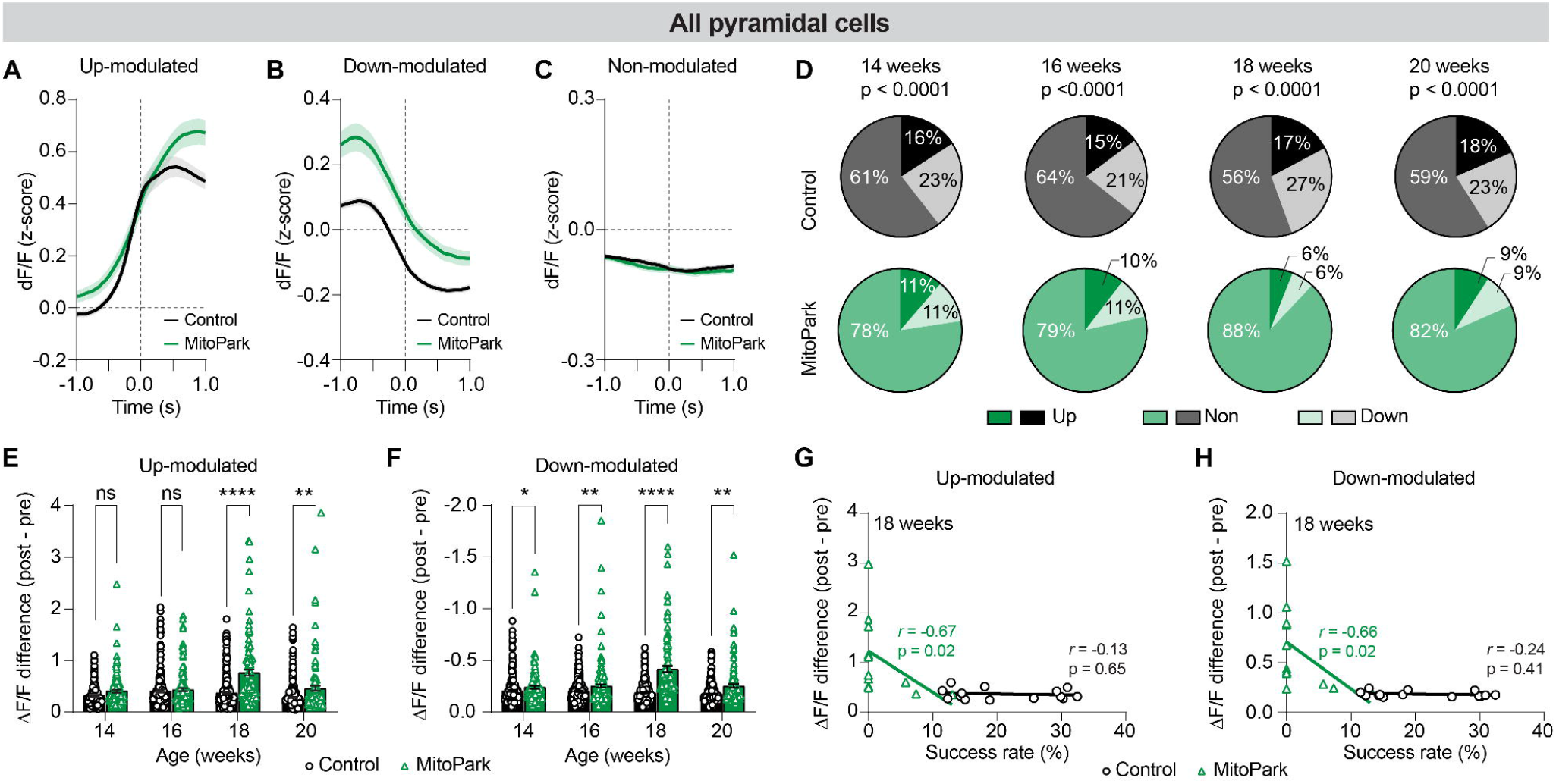
Impaired modulation of pyramidal neurons by movement in Parkinsonism. A-C) Representative mean calcium activity of M1 pyramidal neurons in MitoPark and control mice at 16 weeks of age, showing up- (A), down- (B), and non-modulated (C) categories. Gray dashed vertical lines indicate the grasp onset. **D)** Proportions of different categories of pyramidal neurons at each time point. Chi-square test to compare differences. Control n/N = 1225 cells/11 mice, 1394 cells/13 mice, 1725 cells/14 mice and 1638 cells/14 mice, MitoPark n/N = 1079 cells/13 mice, 1465 cells/12 mice, 1971 cells/12 mice and 1347 cells/9 mice, at 14, 16, 18 and 20 weeks, respectively. **E-F)** Summary results of the magnitude of modulation for up-modulated (E) and down-modulated (F) pyramidal neurons. Linear mixed-effects models for log-transformed data were used (See Methods for details). Each data point represents one cell. (E) Control n/N = 194 cells/11 mice, 206 cells/13 mice, n/N = 297 cells/14 mice and 303 cells/14 mice, MitoPark n/N = 122 cells/13 mice, 152 cells/12 mice, 118 cells/12 mice and 123 cells/9 mice, (F) Control n/N = 289 cells/11 mice, 291 cells/13 mice, 469 cells/14 mice and 369 cells/14 mice, MitoPark n/N = 122 cells/13 mice, 162 cells/12 mice, 119 cells/12 mice and 125 cells/9 mice, at 14, 16, 18 and 20 weeks, respectively. **G-H)** Correlation between success rate and modulation magnitude at 18 weeks, for up-modulated (G) and down-modulated (H) cells, averaged within each mouse. Each data point represents one mouse. Spearman correlation. Control = 14 mice, MitoPark = 12 mice.

Next, we quantified the magnitude of movement-related modulation in pyramidal neurons, i.e., the difference in GCaMP6f fluorescence between pre- and post-grasp onset. We hypothesized that M1 physiological activity pattern correlates with behavioral outcomes. Thus, we compared M1 neural activity changes between successful and failed reaches. In both control and MitoPark mice, successful reaches were associated with a greater magnitude of M1 neural activity changes than failed reaches (Fig. S6). This behavioral outcome-dependent differences were detected across age groups in both up- and down-modulated M1 pyramidal neurons (Fig. S6). These results suggest that the temporal patterns of modulation are more specific to successful reaches than failed reaches. Interestingly, in the up-modulated group, MitoPark mice showed greater increases in the magnitude of GCaMP6f signals than littermate controls at 18 and 20 weeks of age (Fig. 5E). Similarly, in the down-modulated group, MitoPark mice exhibited greater decreases in the magnitude of GCaMP6f signals than littermate controls across ages (Fig. 5F).

We further tested if altered M1 neuronal activities correlate with the severity of impaired behavioral performance of MitoPark mice at 18 weeks of age. We chose this age group because the MitoPark mice showed moderate parkinsonian motor deficits but were able to perform fair numbers of successful reaches. In MitoPark mice, we found a strong and significant negative correlation between the magnitude of neuronal activity changes in both up- and down-modulated groups and the success rate of individual animals (green lines in Fig. 5G, H). However, these correlations were absent in control mice (black lines in Fig. 5G, H). Together these data suggest that MitoPark mice with more pronounced M1 neuronal activity changes showed greater impairment in skilled movements.

Furthermore, we attempted to determine if CSp and IT neurons show different peri-event activity changes in MitoPark mice and littermate controls. Both CSp and IT neurons were categorized into up-, down-, or non-modulated groups (Fig. S7A, B). In control animals, ∼50% of CSp neurons were classified as non-modulated (Fig. S7A). This result is consistent with earlier reports showing that CSp exhibit a heterogeneous relationship between their activity and movement (*45*). As Parkinsonism developed, the proportion of non-modulated CSp neurons gradually increased in MitoPark mice from the early to late stages, and this increase was significantly higher than in littermate controls (Fig. S7A). In both control and MitoPark mice, majority of IT neurons (> 70%) were classified as nonmodulated group, i.e. lacking significant activity changes around grasp onset. These differences in regulating reaching task between CSp and IT neurons might relate to their distinct roles in encoding various aspects of kinematic parameters (e.g., amplitude versus directions) of skilled movements (*43*).

We quantified the amplitude of modulation of neuronal activity in up- and down-modulated CSp and IT neurons, but found no difference between MitoPark mice and littermate controls for either population (Fig. S7C-F). These results were partially due to a significant decline in the number of reaches in parkinsonism, rendering the dataset underpowered to reveal statistical differences. Together, the above results suggest that M1 pyramidal neurons showed progressively impaired movement-modulated activity during the progressive Parkinsonism.

### Longitudinal stability of movement encoding by individual neurons in M1

Leveraging the spatial cellular resolution and stability of miniscope imaging, we followed a subset of CSp neurons from 14 to 20 weeks of age in both control and MitoPark mice, allowing us to study movement-encoding by their activity over time (Fig. 6A). Although the relationship between CSp activity and movement is dynamic and can be altered over time, as reported previously (*45, 47, 51*), a small number of CSp neurons maintained their movement modulation categories over time (Fig. 6A, B), perhaps representing an ensemble of cells important for the analyzed movements (*42*). In control mice, 4% and 10% of up- and down-modulated CSp neurons, respectively, maintained their movement-modulation categories across ≥3 age groups between 14 and 20 weeks (Fig. 6B, C). In contrast, none of the up- or down-modulated CSp neurons could be repeatedly categorized into the same movement modulation category across ≥ 3 ages between 14 and 20 weeks in MitoPark mice (Fig. 6D). These data indicate that the response stability of movement-related activities of individual CSp neurons was gradually impaired in MitoPark mice relative to controls. Quantitatively, we calculated the Fleiss Kappa to assess the stability of movement-modulation by CSp neurons. We found a Fleiss Kappa of 0.27 for CSp neurons in the control group between 14 and 20 weeks, i.e., weak but detectable temporal stability. In MitoPark mice, the Fleiss kappa for CSp neurons was 0.1 between 14 and 20 weeks, indicating no reliable temporal stability. These results suggest a disrupted stability of movement encoding by M1 pyramidal neurons during the progression of parkinsonism (*28*).

**Figure 6.**
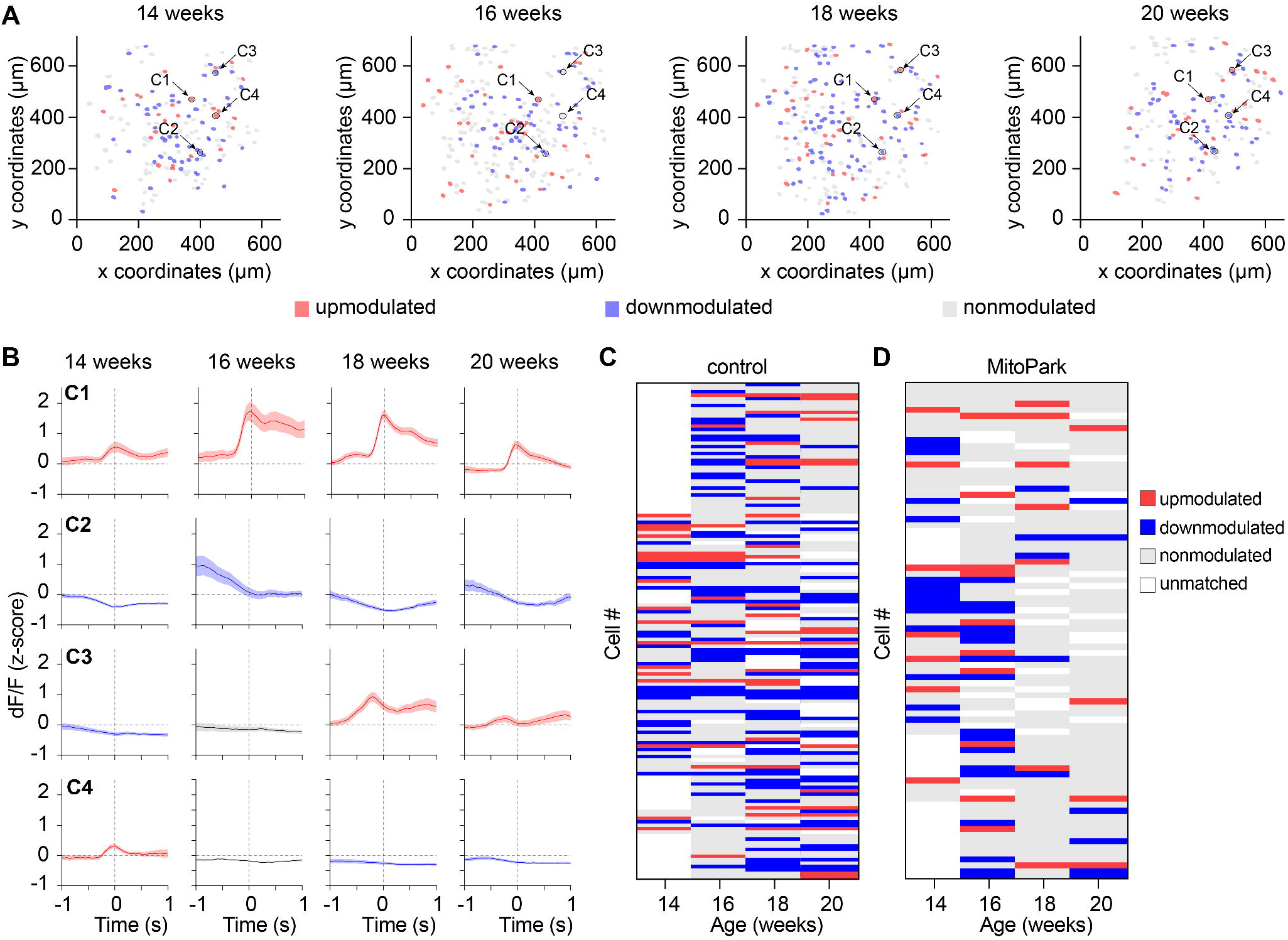
Longitudinal stability of movement encoding by individual CSp neurons in M1. A-B) Representative cell maps (A) and modulation traces (B) from one control mouse, with color coding indicating up-, down-, and non-modulated cells, in red, blue, and gray, respectively. Circled cells in A are the longitudinally registered CSp cells represented in B, showing the mean modulation across trials at different timepoints. C1 and C2 were up- and down-modulated, respectively, across time (stable), whereas C3 and C4 modulation changed across time (unstable). Gray dashed vertical lines indicate the grasp onset. **C-D)** Heatmaps showing modulation of CSp cells registered in at least 3 timepoints, with the same color coding as A and B, for control (C) and MitoPark (D) mice. Each line represents one cell and white indicates missing matching cells. (C) Control n/N = 72 cells/4 mice, 121 cells/6 mice, 102 cells/6 mice and 31 cells/4 mice, (D) MitoPark n/N = 44 cells/3 mice, 59 cells/3 mice, 102 cells/3 mice, and 31 cells/3 mice, at 14, 16, 18, and 20 weeks, respectively.

### Population encoding of movement by M1 neurons is impaired in Parkinsonism

Given the decreased proportion of M1 neurons responsive to movement and their instability in encoding the reach events over time, we sought to understand how these changes affect the dynamic control of skilled movement at the population level. We characterized the temporal dynamics of the denoised Ca^2+^ signals from all GCaMP6f-expressing cortical pyramidal neurons during reaches using an unsupervised spectral clustering algorithm (*52, 53*). Based on their distinct temporal dynamics, cortical pyramidal neurons can be categorized into 7 functional subpopulations (referred to as clusters hereinafter) in both control and MitoPark mice (Fig. 7A-E). These clusters were well separated in a high-dimensional principal component space (Fig. S8A-C) and displayed peak activity at different timings during reaches (Fig.S8D). This analysis suggests the existence of functionally distinct clusters of M1 pyramidal neurons in relationship to the generation of skilled movements. Interestingly, while clusters 2 and 3 overlapped largely with the activity-based up-modulated category, clusters 1, 6, and 7 overlapped with the down-modulated category. All clusters partially overlapped with the non-modulated category (Fig. S8E).

**Figure 7.**
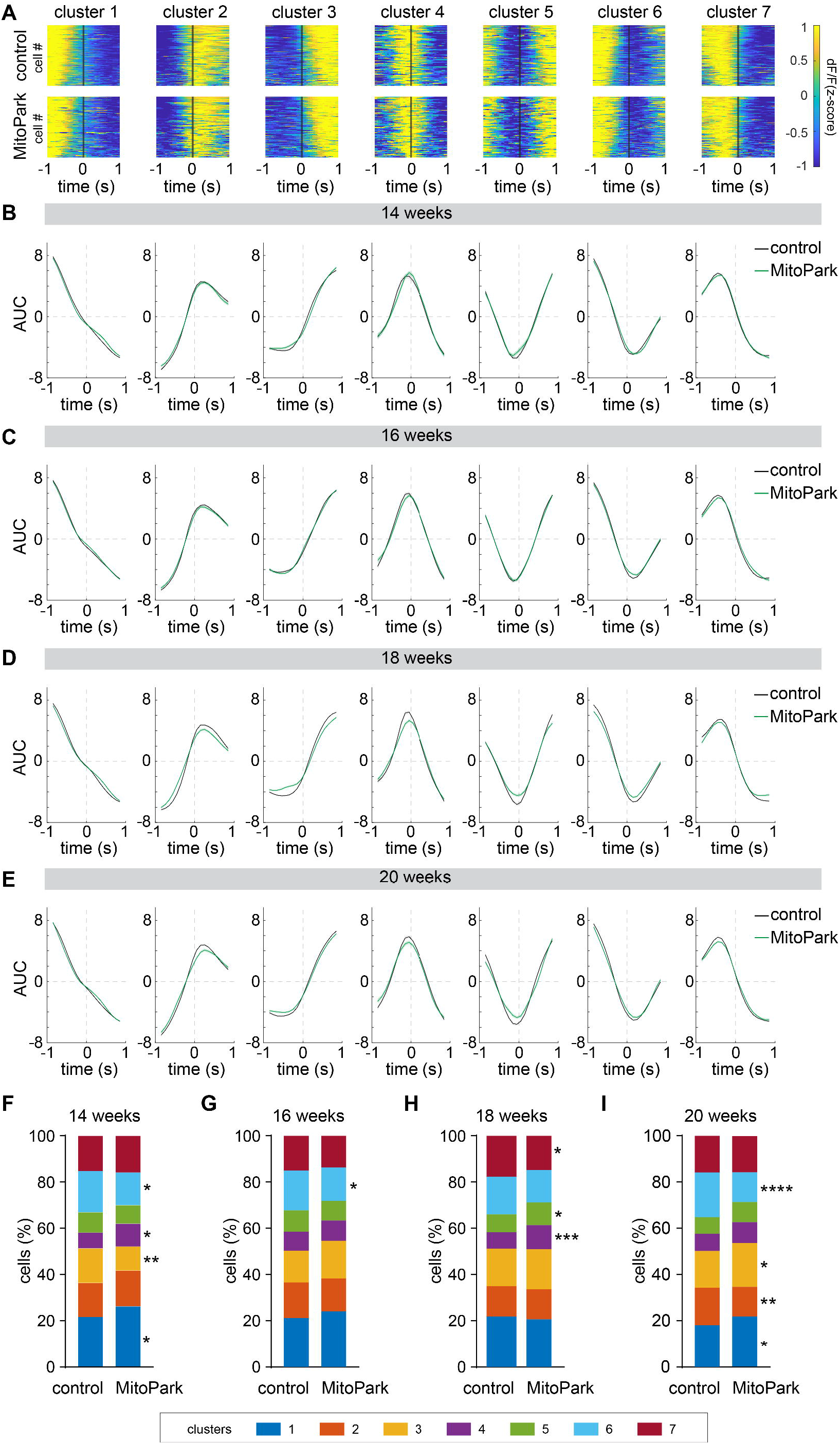
Altered movement encoding by functional clusters of M1 pyramidal neurons in MitoPark mice relative to controls. **A)** Heatmaps of ΔF/F of the different functional clusters of M1 pyramidal cells, aligned to the grasp onset (black vertical line), in control (top) and MitoPark (bottom) mice. Each horizontal line represents one cell. **B-E**) Area under the curve (AUC) of ΔF/F calculated within ±1 sec aligned to grasp onset (gray vertical line). (**F-I**) Cell proportions per cluster in control vs MitoPark mice across ages. The Chi-square test was used to compare groups.

These data support the notion that distinguishable dynamic temporal profiles of movement-related activity of pyramidal neurons exist. Interactions between these subpopulations of M1 pyramidal neurons may contribute to the generation of reaching activity. Thus, we quantified the proportions of cells in each cluster and found significant differences between MitoPark mice and littermate controls across ages (Fig. 7F-I). In particular, the proportion of pyramidal neurons across different clusters remained relatively consistent in control mice of different ages. In contrast, MitoPark mice exhibited longitudinal fluctuations in these proportions between 14 and 20 weeks, resulting in significant group differences in 4 of 7 clusters at 20 weeks (Fig. 7F-I). These results suggest that as parkinsonism develops, movement-related activity patterns across functional clusters of M1 pyramidal neurons are strongly altered, in line with the reduced overall stability of movement encoding in the activity-based categories of neurons in MitoPark mice (Fig. 6).

Next, we implemented binomial logistic regression to assess the contributions of the different clusters to reach decoding. We ran the regression analysis on time-resolved Ca² data from reach epochs and randomly sampled non-reach epochs with matched duration (Materials & Methods). This model reliably distinguished reach from non-reach segments across mice and ages (Fig. 8A), as indicated by consistently high balanced accuracy (a metric that evaluates how well the model identifies reach/non-reach segments). There was no correlation between the balanced accuracy and trial or cell numbers (Fig. S9). Classifier performance was further evaluated by examining the variability of model weights across folds, which was minimal, and by inspecting confusion matrices, which consistently showed high fractions of correct predictions (Fig. 8B).

**Figure 8.**
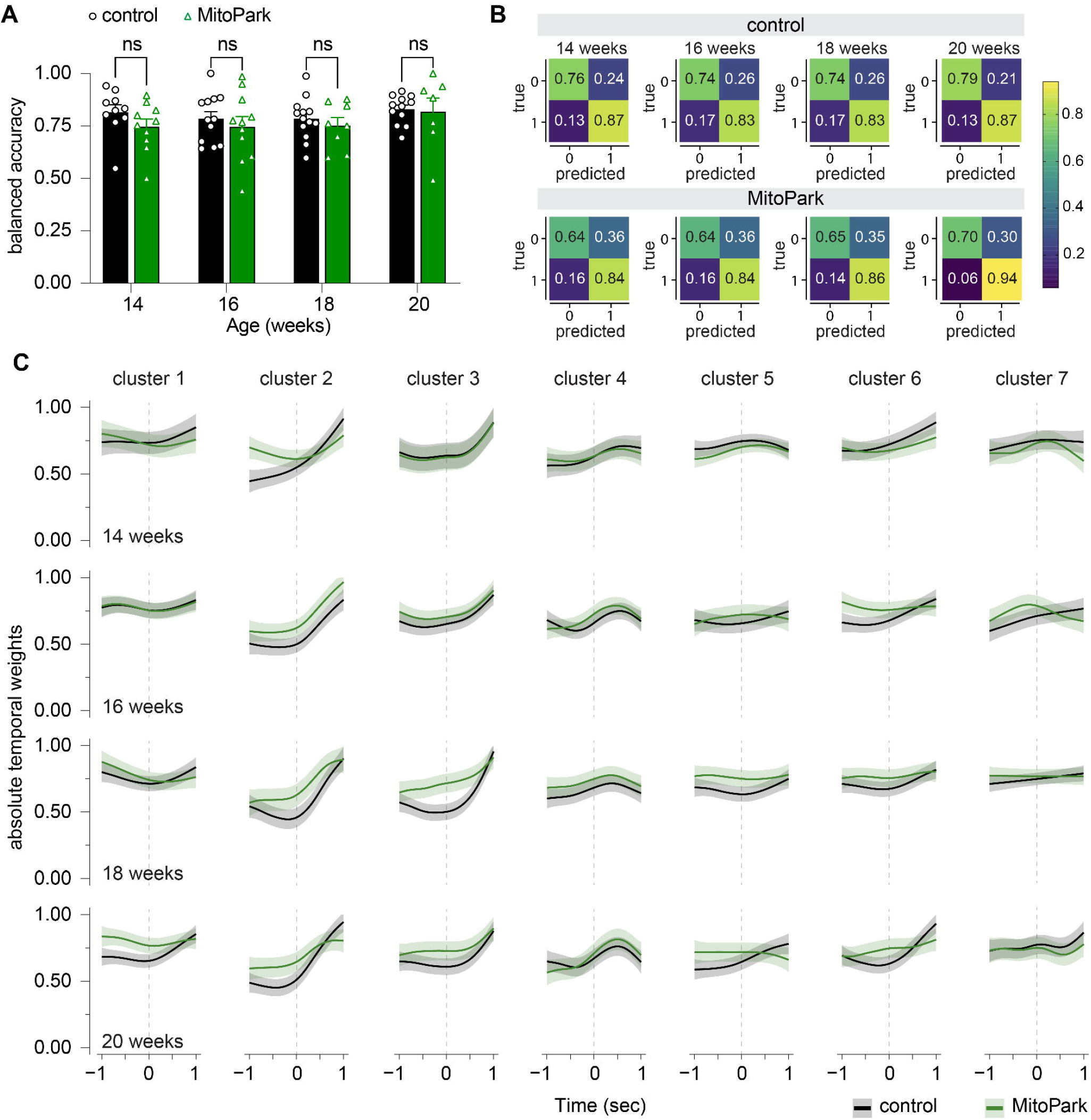
Predictive modeling indicates an abnormal cortical neuronal encoding of movement in progressive Parkinsonism. **A)** No difference was observed between MitoPark and controls in the balanced accuracy of the binomial logistic regression model, averaged across folds. Linear mixed-effects model (see Methods for details). Each data point represents one mouse. **B)** Confusion matrices averaged across PCA folds for control (top) and MitoPark mice (bottom) groups, reported as fractions. (A-B) Control N = 10, 12, 13 and 12 mice, MitoPark N = 10, 12, 8 mice and 7 mice, at 14, 16, 18 and 20 weeks, respectively. **C)** Absolute temporal weights of the different clusters from the logistic regression model in control and MitoPark mice across ages. GAMM with autocorrelated errors was fitted for each functional cluster at each age, with separate smooths for time in MitoPark and control. Shown are the fitted smooth curves and point-wise 95% confidence bands. The difference curves are shown in the supplementary Fig. 10.

Furthermore, we explored possible changes in the contributions of different clusters to movement decoding by comparing the absolute model weights across ages (as a measure of each cluster’s contribution to the model’s outcome) in function of time around grasp onset (Fig. 8C). We found divergence in clusters’ contribution to movement decoding between MitoPark mice and littermate controls, particularly at 18 and 20 weeks of age (Fig. 8C, Fig. S10). Together, these data demonstrate the altered stability of movement-related activity of M1 pyramidal neurons at both individual neuron and populational levels in progressive Parkinsonism (*28*).

## DISCUSSION

In the present work, we studied neuronal activity changes in M1 and examined their relationship to impairment of skilled movement during the progression of Parkinsonism (Fig. 9). We can draw a conclusion that MitoPark mice showed gradual impairments in motor skills associated with possibly disrupted kinematic control of forepaw movements as they develop Parkinsonism (Figs. 1 & S2). The impaired skilled movements were accompanied by decreased activity of nonspecific pyramidal neurons in M1 (Fig. 2). At the cell-subtype level, movement-related activity of CSp neurons is preferentially reduced over time in MitoPark mice (Fig. 3). These observations support the conclusion that M1 circuits develop cell subtype-specific alterations in Parkinsonism (*18, 19, 22, 23*). Moreover, MitoPark mice exhibited fewer co-activated M1 pyramidal neurons as they developed Parkinsonism (Fig. 4), which was accompanied by greater movement-related activity changes than in controls (Fig. 5). It remains to be defined whether the greater movement modulation of single neuron activity is a compensation for the fewer coordinated activation of M1 pyramidal neurons at population level, or they are two independent aspects of cortical pathophysiology in PD. Furthermore, at the population level, M1 pyramidal neurons can be categorized into distinct functional clusters based on their temporal profiles relative to movements (Fig. 7). The dynamic contribution of these clusters to movement decoding appears to differ gradually and notably between MitoPark mice and controls over time (Fig. 8). Together, the present study reveals an abnormal and unstable relationship between M1 neuronal activity and the generation of skilled movement under Parkinsonian states (*21*).

**Figure 9.**
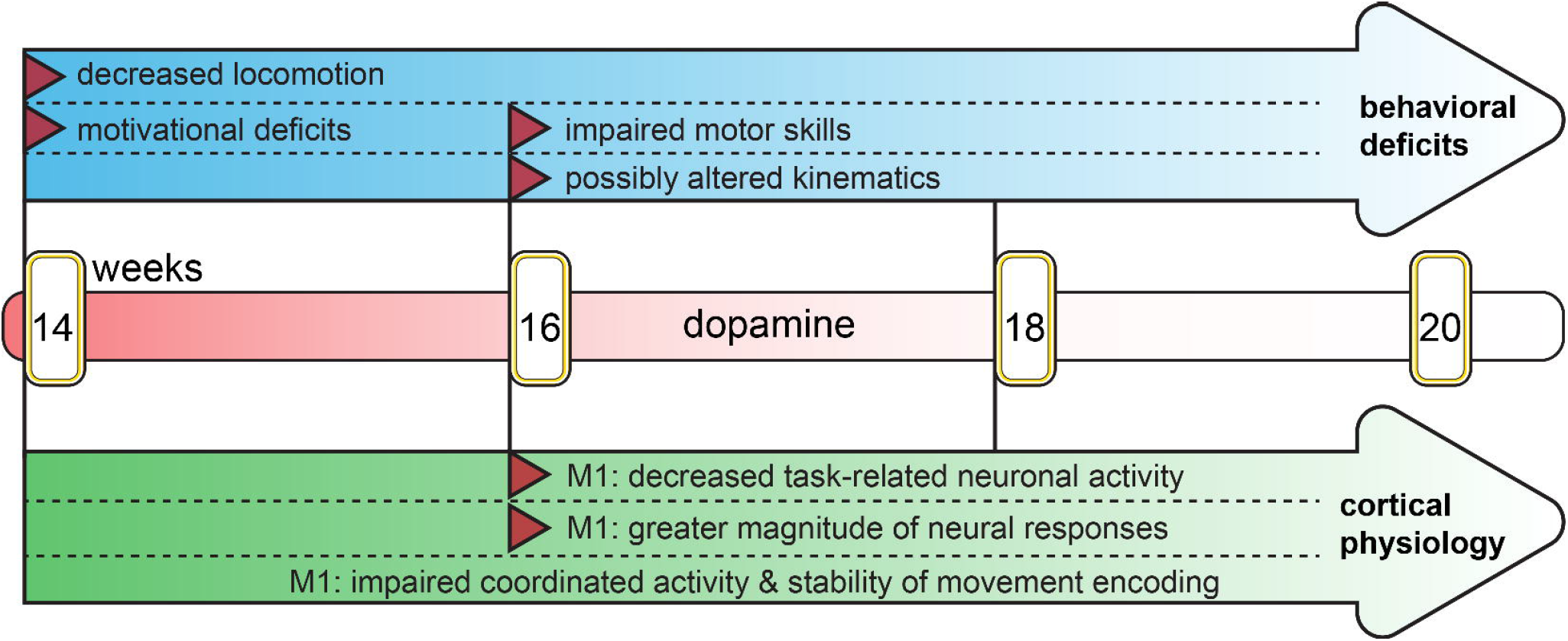
Graphical abstract. Diagram highlighting the temporal profile of behavioral deficits (top) and the emergence of cortical pathophysiology (bottom) during progressive loss of dopamine (middle) in MitoPark mice.

### Cell-subtype-specific M1 circuit changes in Parkinsonism

Although the traditional circuit model of PD pathophysiology predicts that M1 is hypoactive due to abnormal basal ganglia inhibition following midbrain dopamine neuron loss, emerging evidence suggests that M1 circuits develop cell-subtype- and input-specific adaptations in Parkinsonism (*18, 19, 22, 23*). Here, we provide additional information on the time course of M1 circuit changes in freely moving MitoPark mice engaged in movements. We confirm that CSp neurons exhibit greater task-related activity changes *in vivo* than IT neurons. Thus, both *in vivo* and *ex vivo* electrophysiological and Ca^2+^ imaging studies demonstrate that the Parkinsonism-related neuronal dysfunction in M1 differs by cell type (*18, 19, 22*). In addition, we found that CSp neurons are hypoactive in 16-week-old MitoPark mice with nearly complete striatal DA depletion and moderate impairment of general motor function (*31*). Notably, CSp neural activity changes during reaching epochs occurred slightly later than that of nonspecific M1 pyramidal neurons that started at 16 weeks (Figs. 2 & 3). In addition, while nonspecific M1 pyramidal neurons in MitoPark mice showed reduced Ca^2+^ event frequency during non-reaching epochs (Fig. 2), such reduction was absent from CSp and IT neuronal dataset (Fig. 3). These discrepancies might be at least partially due to the biased cell labeling associated with different viral vectors, or underpowered experimental design.

### Altered movement modulation by M1 pyramidal neurons in Parkinsonism

Layer 5 CSp neurons directly connect the M1 circuit with subcortical motor regions and critically contribute to the execution of movement (*45, 54*), but they are relatively less studied using optical imaging approaches compared to layer 2/3 neurons (*42, 46, 55*). This is at least partially due to their deep location, making them less accessible with *in vivo* two-photon imaging. By implanting a prism GRIN lens, we were able to monitor deep CSp neuronal activity in freely moving control and Parkinsonian mice for months. In control mice, we found noticeable heterogeneity in the relationship between the activity of CSp neurons and movement. Specifically, the activities of ∼50% of CSp neurons showed no correlation with the monitored skilled movement, whereas the activities of the remaining proportion of CSp neurons correlate positively or negatively with movement. Such heterogeneity in the correlation between CSp activity and movement has also been reported in prior research (*45, 56*). It is noteworthy that the activity of a large proportion of CSp neurons is modulated by well-learned skilled movements in control mice, indicating their role in motor execution. This is distinct from the gradual disengagement of L2/3 pyramidal neurons from learned movement over time (*55*). During progressive Parkinsonism, the proportion of movement-modulated cortical pyramidal cells gradually decreased. In addition, these neurons also showed a less stable temporal relationship to movements in MitoPark mice than in controls. Furthermore, the proportions of cortical pyramidal neurons within distinct functional clusters (Fig. 7) were significantly altered in MitoPark mice compared with littermate controls. Together with the longitudinal tracking of a subset of cortical pyramidal neurons (Fig. 6), these data suggest a reduced stability of the movement-related cortical neuronal activity during progressive parkinsonism. Combined with the changes in spine dynamics of and synaptic inputs to pyramidal neurons (*19, 28, 34, 40, 42*), as well as other functional impairments (*33, 47, 57–60*), these M1 microcircuit adaptations during the progression of Parkinsonism may be important determinants of the cardinal motor signs of PD.

### Molecular, cellular and circuit mechanisms of M1 dysfunction in parkinsonism

This work described numerous behavioral and cortical neuron activity changes during progressive Parkinsonism (Fig. 9), which could involve several molecular and cellular mechanisms. In healthy controls, acquisition of skilled movements is associated with coordinated activity of M1 neurons (i.e., functional ensembles), a process involving synaptic plasticity (*42, 61*). In PD, striatal DA depletion triggers cellular and synaptic adaptations throughout the basal ganglia nuclei and the thalamocortical network. Specifically, degeneration of midbrain DA neurons results in loss of dendritic spines and downregulation of the intrinsic excitability of M1 pyramidal tract neurons in Parkinsonian NHPs and rodents (*18, 19, 24*). The weakened thalamic excitatory inputs and loss of dendritic spines are likely to break down the thalamocortical network (*19, 24*), underlying less coordinated and more independent M1 neuronal activity during movement (Fig. 4). Moreover, our recent work reported that the thalamocortical synaptic deficits occur in 16 weeks old MitoPark mice (*31*). Considering the temporal development of motor deficits and M1 network dysfunction, the impaired thalamocortical connectivity could at least partially explain the reduced co-activated M1 pyramidal neurons associated with skilled movement (Fig. 4) and smaller proportion of movement-modulated pyramidal neurons (Fig. 5D) in MitoPark mice relative to controls. While fewer coordinated pyramidal neurons were engaged in skilled movement, the movement-modulated neurons exhibited greater magnitude of response (Fig. 5E, F). This observation indicates the involvement of unidentified M1 local microcircuits that enhance cellular responsiveness to compensate for the reduced synaptic connectivity of M1 pyramidal neurons, including maladaptive changes of the other glutamatergic inputs or GABAergic interneurons (*31*).

Additionally, the altered movement encoding by functional clusters of M1 pyramidal neurons likely involves intricate interactions among synaptic excitation and inhibition as well as cellular excitability (*56, 62*). In the present study, animals underwent extended training to perform uncued skilled movements. It has been reported that inactivation of premotor regions impairs the performance of such internally-generated skilled movement (*60*). Our unpublished observation indicates that cortical input from the secondary motor cortex to M1 decreases significantly following DA loss. Thus, while the worsened motor performance is accompanied by decreased pyramidal neuronal activity, other impairments, such as changes in corticocortical interactions, may contribute to Parkinsonian motor signs.

Furthermore, dopaminergic projections from the ventral tegmental area (VTA) can modulate M1 neuronal activities and circuit function, contributing to the acquisition and execution of skilled movements (*37, 63–65*). Prior work reported that local dopaminergic signaling results in altered dynamics of dendritic spines and synaptic plasticity (*40*) as well as impaired cellular excitability and thalamocortical inputs (*63, 66*). The mesocortical dopaminergic projections and VTA DA neurons could be affected in MitoPark mice at 20 weeks and beyond, leading to at least a partial DA depletion in M1 (*29, 35*). Therefore, the involvement of local cortical dopaminergic neuromodulation in the observed behavioral and cortical physiological changes of MitoPark mice can’t be fully excluded.

### Limitation of the present work

While longitudinal *in vivo* recordings from freely moving mice are a technical strength of the present work (*67*), freely moving mice initiate and execute internally generated reach-to-grasp behavior at a self-defined pace, making data analysis challenging. One limitation of the present work is the reduced number of reaches as Parkinsonism progresses in MitoPark mice, making it impossible to study the relationship between M1 activity and different reach categories (e.g., successful v.s. failed) due to the small sample size, particularly in animals with advanced Parkinsonism. A limitation of the Ca^2+^ imaging approach is its low temporal resolution does not match the rapid reaching activities in mice. Such a mismatch in temporal dynamic makes detailed investigation of the relationship between individual neuronal activity and movement parameters impossible.

The slow kinetics of GCaMP6f signals is not appropriate for testing high-frequency oscillatory neuronal activity. However, as shown in this study, Ca^2+^ imaging provides valuable information that advances our understanding of PD pathophysiology, especially in longitudinal studies of defined groups of neurons, which may help guide the design of effective cortex-based therapies for the disease. Furthermore, the progressive and robust neurodegeneration, as well as adult-onset parkinsonism are the strength of the MitoPark mice compared to neurotoxin-based or genetic models of PD (*68*). We would like to point out that this model does not develop α-synuclein protein aggregation that is a featured neuropathology in human PD (*29*). In addition, MitoPark mice develop mainly dopaminergic neuropathology, but do not affect other vulnerable neuromodulatory systems that degenerate in PD, like the noradrenergic neurons in the locus coeruleus (*29, 35*). That said, additional neural mechanisms are expected to contribute to cortical pathophysiology of the impaired motor skills in PD patients.

## MATERIALS AND METHODS

All animal studies were reviewed and approved by the Georgetown University Institutional Animal Care and Use Committee (IACUC protocol #2024-0001).

### Animals

DAT^Cre^ mice (JAX stock #006660) were crossed with Tfam^fl/fl^ mice (JAX stock #026123) to obtain double heterozygous mice (DAT^Cre/+^, Tfam^fl/+^), which were then crossed with homozygous Tfam^fl/fl^ mice to obtain MitoPark mice (DAT^cre/+^, Tfam^fl/fl^) for experiments (*29, 30*). Mice were housed up to 5 animals/cage under a 12h/12h light/dark cycle with free access to food and water in accordance with NIH guidelines for animal care and use.

### Stereotaxic surgery

For *in vivo* cell-subtype visualization, we performed stereotaxic surgery at 5-10 weeks of age to retrogradely label subtypes of M1 pyramidal neurons based on their long-range axonal projections. Specifically, mice were placed on a stereotaxic frame (RWD, Model: 71000) under 2-3% isoflurane for anesthesia and were subcutaneously injected with carprofen (5 mg/kg) or buprenorphine (3.25 mg/kg) for analgesia. Body temperature was maintained and monitored using a heating pad connected to a feedback controller. To record calcium activity from cortical excitatory neurons, AAV9-CaMKII-GCaMP6f (Addgene #100834-AAV9, RRID: Addgene_100834-AAV9; volume = 300 nl, titer = 1 × 10^13^ GC/ml) was injected in M1 contralateral to the dominant paw [from bregma in mm, anteroposterior (AP) = +0.3 and +0.8, mediolateral (ML) = 1.25, dorsoventral (DV) = −1.0 from dura]. To retrogradely label IT neurons, AAVrg-CAG-tdtomato (Addgene #59462-AAVrg, RRID: Addgene_59462-AAVrg; volume = 500 nl, titer = 7 × 10^12^ GC/ml) was injected in the dorsolateral striatum (DLS) that was contralateral to the M1 injections (in mm, AP = +0.7, ML = 2.3, DV = −2.6, and AP = +1.0, ML = 2.0, DV = −2.6 from dura, 500 nl per injection). Details of intracranial injection surgery can be found on Protocols.io (10.17504/protocols.io.rm7vzye28lx1/v1). To retrogradely label corticospinal neurons in M1, AAVrg-CAG-tdtomato (Addgene #59462-AAVrg, RRID: Addgene_59462-AAVrg; titer = 7 × 10^12^ GC/ml) was injected in spinal C7-8 segments (ML = ±0.5 from midline and DV = −0.7 from dura). Details of spinal cord surgery can be found on Protocol.io: 10.17504/protocols.io.81wgbz5e3gpk/v1. Mice used for IT neuron studies were trained to perform the SPRG task between 5 and 8 weeks of age (see below), followed by AAV injections (Fig. 1A). Considering the potential spinal cord damage associated with invasive surgery that may impair the acquisition of motor skills, mice for CSp neuron studies were subject to AAV injections into the spinal cord at 5 weeks of age, and then allowed to recover for 2 weeks before motor training between 7 and 10 weeks of age (Fig. 1A). Once mice acquired the SPRG skills, they received AAV9-CaMKII-GCaMP6f injections and Prism lenses implantations in M1. After the surgery, all mice were retrained for 1-2 weeks to ensure that they could perform the SPRG task.

### GRIN lens implantation

Mice were mounted on a stereotaxic frame (RWD, Model: 71000) under 2-3% isoflurane anesthesia and subcutaneously injected with carprofen (5 mg/kg) or buprenorphine (3.25 mg/kg) for analgesia. Body temperature was maintained and monitored using a heating pad connected to a feedback controller. A prism GRIN lens attached to a baseplate (Cat # 1050-004419, Inscopix) was inserted in M1 at the AAV injection site centered at (in mm) AP = +0.5, ML = 1.0, DV = −0.5, so that the imaging plane would be parallel to the midline and faces M1 medially. The lens was held using a dummy microscope (Inscopix, Cat # 1050-003762) and lowered at 2 μm/min. The lens and baseplate were fixed to the skull using C&B Metabond dental cement (Parkell, Farmingdale, NY), and a baseplate cover (Inscopix, Cat # 1050-004639) was placed to protect the lens. Mice with lens implantation were single-housed after surgery. Details of lens implantation surgery can be found here: https://www.protocols.io/view/injection-of-aav-and-prism-grin-lens-implantation-4r3l21rrqg1y/v1.

### Behavior training and data analysis

Details of the construction of the behavior training box and the SPRG task training protocol were described in our methods paper (*32*). Briefly, mice were habituated to the experimenter and the customized training box for 5 days. Then, mice were food-restricted to keep the body weight at 80% of baseline until the end of training. Mice underwent 2 to 3 days of shaping sessions (20 min), during which they learned to use their forepaws rather than their tongues to obtain pellets. The dominant forepaw was identified as the one used for more than 70% of the reaches. Mice that consistently grasped pellets using their paws entered the training phase. This included daily 20-minute sessions over 7 days, during which the mice were trained to use only their dominant paws to grasp the pellets. Forepaw movement was recorded using a high-speed FLIR camera (Teledyne FLIR, Model: Blackfly S) that was triggered by the custom task controller when the animal approached the frontal slit and placed its paws on a metal bar. Behavioral videos were reviewed by an experimenter blinded to the animal genotypes using BORIS software (Behavioral Observation Research Interactive Software, University of Torino, RRID:SCR_025700) to manually identify overall reaches, failed and successful reaches, as well as to timestamp the start of the reach, onset of grasp (i.e., the peak timing of reaches) and the end of the reach. The onset of grasp was used for peri-event reaching analysis, and the start and end were used for reach kinematics analysis. Details of the behavior training protocol can be found here: https://www.protocols.io/view/mouse-reach-to-grab-task-training-protocol-ewov11bb2vr2/v1. To compare all behavioral variables described below (Fig. 1B, C), a linear mixed-effects model with fixed effects for group, age (a factor with four levels), and the group-by-age interaction, with a random intercept for mouse to account for repeated measures was fitted. We then used Kenward-Roger degrees of freedom and Benjamini-Hochberg correction was applied across the age-specific group comparisons.

#### Overall motor skills assessment

To assess skilled movements, we defined four main categories of reaches based on the outcomes: 1) successful reaches: the mouse successfully grasps and retrieve the sugar pellet to its mouth; 2) drop reaches describe when the mouse successfully grasps the pellet, but loses it during the retrieval phase; 3) flick reaches: the mouse touches the pellet but fails to grasp it, flicking it off the reachable position; and 4) miss reaches: the mouse doesn’t even touch the pellet. The last three are considered failed reaches. The rate of each category was computed as its proportion out of the sum of all four categories. For example, success rate = (number of successful reaches)/(the sum of successful, drop, flick, and miss reaches) x 100%. The final category, “in vain” reaches, describe where the mouse continues to perform reaching movements even though a pellet is not present. Thus, total number of reaches = the sum of (successful, failed, and in vain reaches).

#### Motivation and task engagement assessment

To quantify motivation and task engagement, we first calculated total number of reaches. This total includes the 4 categories successful and failed reaches described above, plus in vain reaches. We also compared the latency to start the first reach (extracted from manual timestamps) and the time on task. The time on task was measured by recording the time the mouse spent holding on a metal perch located adjacent the food retrieval slit in the front of the training box (Fig. 1F).

#### Movement kinematics

For a subset of mice (3 per group) we acquired side view videos at 100 fps, by placing a camera on the same side as the reaching paw. A DeepLabCut (version 3.0.2, RRID:SCR_021391) network was trained on 1200 frames to track the wrist joint of the reaching paw, the center of the paw, and the 4 digits (*69*). The wrist joint and the digits were labeled to provide anchor points and build a skeleton thus improving the paw tracking (Fig. S2A-C). The center of the paw was the only point included in subsequent analysis. We used a resnet50 model trained for 200 epochs with default parameters and evaluated manually for the accuracy. This model was then used on all the videos to obtain horizontal (*x*) and vertical (*y*) trajectories. Using MATLAB (version R2026a, Mathworks, RRID: SCR_001622) built-in functions, the obtained *x*,*y* trajectories were converted to mm, with the metal perch as the start point. They were first segmented using the manually timestamped reach start, grasp onset, and reach end to obtain the coordinates for each reach movement separately. We then filtered the paw coordinates such as to only retain frames with a tracking likelihood more or equal to 0.55; frames below this threshold were treated as missing values. Next, outliers were removed using a moving median with a 7 frame window (*isoutlier()*, with method ‘movmedian’). Then, gaps of 3 to 5 consecutive missing frames were interpolated using shape-preserving piecewise cubic spline interpolation (*fillmissing()* with method ‘pchip’). Reaches with longer gaps were excluded from the analysis. The resulting trajectories were smoothed using a third-order Savitzky–Golay filter with a 7 frame window (*smoothdata()* with method ‘sgolay’). To quantify the movement kinematics, we computed the full length of the trajectory and the mean speed of all reaches. Due to the limited number of mice in each group, we pooled the extracted reaches and used a two-way ANOVA followed by Sidak test for group comparisons at each timepoint. Then, we focused on successful and missed reaches at 16 weeks when MitoPark mice developed moderate Parkinsonism but still performed a fair number of successful reaches (Fig. 1 and Fig. S1), and computed the instantaneous speed to find the maximum speed. We resampled speed linearly to 50 points (*interp1()* with method ‘linear’) then expressed timing of max speed as a fraction of the reach duration. To assess how stereotypical the movement was across reaching trials, we computed pairwise similarity between the trajectories (*70*). First, trials were linearly resampled to 50 points (*interp1()* with method ‘linear’). Next, for each trial, the horizontal (*x*) and vertical (*y*) positions of the paw were centered by subtracting their own temporal mean to only preserve the shape of the trajectory. The two vectors were then joined end to end into a single long vector so that both spatial dimensions contributed to a single measure of similarity. For each pair of trials, the similarity was defined as the Pearson correlation coefficient between their vectors. We then quantified the number of pairs of reaches with *r* above and below 0.7 (Fig. S2).

### *In vivo* Ca^2+^ imaging

*In vivo* calcium imaging was performed using nVue 1.0 (Inscopix, Cat # 1000-005735) and nVue 2.0 (Inscopix, Cat # 1000-006810) systems. Two weeks post-surgery, the mice were placed in the behavior box daily for 7 days to re-habituate to the setup with a miniscope attached to the baseplate. At the end of the habituation, the mice were fasted overnight, followed by *in vivo* imaging the next day. During the imaging session, mice were placed in the behavior box with the miniscope attached to the baseplate. While mice were in the training box, the GCaMP6f signal was recorded continuously at 20 Hz using the Inscopix Data Acquisition System by setting LED power at 0.2 -0.4 mW/mm^2^, exposure time at 50 ms, and sensor gain of 2 to 4. During recording, transistor–transistor logic (TTL) pulses (5 V) were generated by the behavioral camera and sent to the nVue system to synchronize GCaMP6f recordings and behavior videos. Digital focus is manually chosen to obtain more cells in the field of view, typically between 0 and 1000 Inscopix units (i.e., ∼ 0-300 μm). The static red channel videos were also recorded for 5 sec at multiple focal planes for cell-subtype identification. At the end of the recording session, the mice were returned to their cages with food *ad libitum*. Details of Ca^2+^ imaging protocol can be found here: 10.17504/protocols.io.yxmvm1w5nv3p/v1

### Ca^2+^ data analysis

All image processing was performed with Inscopix Data Processing Software (IDPS). Spatial downsampling by a factor of 2 was applied to the images for efficient data processing, whereas no temporal downsampling was applied. Videos were motion-corrected, and cell traces were extracted using the CNMF-E algorithm (*71*), which is widely used for background-fluctuation correction of one-photon calcium imaging data. Cell traces and contours extracted by CNMF-E were manually checked and artefacts were discarded. The red channel videos were used to categorize CSp or IT neurons, which were used for further analysis.

#### Ca^2+^ events detection

Ca^2+^ signals were analyzed by behavioral states: reaching versus non-reaching epochs. We first identified non-reach epochs, defined as periods of at least 30 seconds during which the mouse did not perform any reaches. Once these periods were extracted, the remaining epochs were identified as reach epochs. Quantification of Ca^2+^ events from signal traces (e.g., frequency, AUC of individual Ca^2+^ transients) was performed on the IDEAS platform. First, the GCaMP6f traces of the cells identified by the CNMFe were denoised using the Deconvolution workflow on IDEAS, using the median method for estimating pixel noise, with a minimum noise frequency of 0.25 and a maximum noise frequency of 0.5. Then, the Ca^2+^ event detection algorithm was applied to the denoised traces. Briefly, baseline fluctuations were removed by subtracting the median trace using a 200 time bins sliding window and a 5-frame sliding average (*72*). Events were then extracted by searching for local maxima based on the following criteria: the amplitude at the maximum must be more than 2 standard deviations (SD) from the baseline, the width must be at least 10 frames when the mean intensity surrounding the maximum remains above 2 SD, and there must be a separation of at least 10 frames between two consecutive events (*72*). The amplitude of each event was measured at the peak prominence of the event starting from baseline. The area under the curve (AUC) was measured at half-width prominence (*72*). To statistically compare the frequency, amplitude and AUC of Ca^2+^ events, during reach and non-reach epochs, outliers were first identified using median absolute deviation method and removed prior to analysis. GCaMP6f event frequency of all cortical pyramidal neurons (Fig. 2) was then analyzed using linear mixed-effects models (R’s *lmer()* from *lmerTest.*). Group, age, and behavioral state (reaching vs. non-reaching) were entered as categorical fixed effects. To account for the hierarchical and repeated-measures structure of the data, all models included random intercepts for mouse, for age within mouse, for behavioral state within age within mouse (i.e., each recording session), and for individual cells. Cell identifiers were made unique across mice and ages, so that the cell term captured dependence between a cell’s values in the reaching and non-reaching states. We first tested the group × age × behavioral state interaction by comparing the full model with a model lacking this term. Because the three-way interaction was not significant (p = 0.75), we then fit a model containing all two-way interactions and tested the group × state interaction by comparing it with a model lacking that term. Model comparisons used F-tests with the Kenward-Roger approximation for denominator degrees of freedom followed by Benjamini–Hochberg correction. Within each behavioral state, another model was fit for frequency, with group, age, and their interaction as fixed effects, with mouse as the repeated measure random effect and an additional random intercept for each mouse-at-age combination (Fig. 2F-G). This latter model was also used for amplitude and AUC (Fig. S4 and S5). The significance of fixed effects was assessed using analysis of variance with Kenward-Roger’s degrees of freedom followed by Benjamini–Hochberg correction. To compare differences in Ca^2+^ events by cell subtypes (i.e., CSp versus IT neurons) under different behavioral states, a linear mixed-effects model with fixed effects for cell subtype, age, group, their pairwise interactions, and their three-way interaction, with random intercepts for mouse and for age nested within mouse to account for repeated measurements was fit (Fig. 3C, D, G, H). Within each model, the Benjamini-Hochberg correction was applied across the age-specific group comparisons.

#### Coactivation analysis

To assess pairwise coactivation of calcium signal, we quantified the temporal co-incidence of detected calcium-event onsets between cell pairs. For both MitoPark mice and controls, we randomly sampled 45-second segments of GCaMP6f recordings from nonspecific pyramidal neurons under reach and non-reach epochs for this analysis. The duration of 45-sec segments was chosen based on the balance between having meaningful recordings (i.e., number of Ca^2+^ events) and including enough animals for proper statistical power, considering that MitoPark mice developed gradual declines in the number and duration of reach epochs. Calcium-event onsets were obtained from the event detection pipeline. Two event onsets were considered co-incident when they occurred within ±100 ms (±2 frames) of one another. For each cell pair, the observed coactivation rate was normalized by the number of events in the less active neuron. To generate a null distribution, the event-onset train of one neuron was circularly shifted relative to the other neuron, thereby preserving each neuron’s event rate and temporal structure, while disrupting their temporal alignment. Coactivation was calculated across 1,000 randomly selected circular shifts. Pairwise coactivity was expressed as a Z-score relative to the shuffled distribution. The co-activation Z-score was calculated as the difference between the observed co-activation and the mean co-activation across circularly shifted data, divided by the standard deviation of the shuffled distribution. We then calculated for each mouse the proportion of cell pairs with Z ≥ 2, corresponding to observed co-activation that was at least two standard deviations above the circular-shift null distribution. For group comparisons across age, we fit a linear mixed-effects model with fixed effects for group, age, and the group-by-age interaction, with a random intercept for mouse to account for repeated measures. We then used Kenward-Roger degrees of freedom and Benjamini-Hochberg correction.

#### Peri-event Analysis

Reach-related Ca^2+^ activity was performed using the peri-event Analysis workflow on the Inscopix IDEAS platform. This workflow quantifies changes in neural signals surrounding event timestamps, i.e., the endpoint of reaches, for each session. This is done at both the population level and for individual neurons, categorizing them as up-modulated, down-modulated, or non-modulated. To do so, we isolated bouts of reaches defined as those not preceded by another reach within at least a 2-second interval. Then the denoised Ca^2+^ traces and the timestamps of the first reach in a bout were fed to the peri-event analysis workflow, which outputs the 3 modulation categories, compared to a null distribution obtained from 1000 random shuffles of event time. The statistical window was set to ±1 sec around the onset of grasps, with a significance threshold of 0.05.

For up- and down-modulated cells, we quantify the magnitude of modulation by measuring the difference in the ΔF/F values between the +1 and -1 sec time windows. For group comparisons across ages (Fig. 5E-F), the up-modulated ΔF/F differences were log-transformed, and the down-modulated ΔF/F differences were transformed with -log(-ΔF/F difference). Then, linear mixed-effects models for log-transformed data were fitted with fixed effects for group, age, and their interaction, and random intercepts for mouse and age, with age nested within mouse. The random intercept for mouse captures correlation between cells on the same mouse and the random intercept for age nested in mouse captures additional correlation from cells recorded in the same week. Within each model, the Benjamini-Hochberg correction was used for multiple comparisons. To further assess correlation between motor impairment and modulation magnitude, we ran a Spearman correlation (using MATLAB’s *corr()* function with ‘spearman’ method) on the success rate and ΔF/F differences for both up and down-modulated neurons of each experimental group at 18 weeks, when MitoPark mice developed moderate Parkinsonism and performed reasonable numbers of successful reaches (Fig. 5G-H).

#### Longitudinal registration

The *CaImAn* Multi-Registration tool on the Inscopix IDEAS platform was used to track cells across sessions. This tool is built around the main function *register_multisession* from the registration algorithm implemented in *CaImAn* (*73*). The workflow was run across multiple sessions for each mouse, with x/y pixel shifts ranging from 10 to 50 pixels, 10 pixels maximum distance between matching centroids, and a maximum footprint disjunction of 0.7. The matched cells were used to compare cell modulation stability over time between groups, based on results from Peri-Event Analysis (matching cells across at least 3 sessions). To assess the stability of movement encoding by CSp neurons, agreement in categorical classifications across time was quantified using chance-corrected kappa statistics in a custom MATLAB script. For pairwise comparisons between consecutive timepoints, Cohen’s κ was computed as 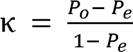 where *P_0_*is the observed proportion of agreement and *Pe* is the expected agreement by chance. Overall agreement across all timepoints was assessed using Fleiss’ κ, defined as 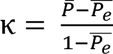 where *̄P* is the mean observed agreement across subjects and *̄P_e_* is the expected agreement based on category marginal proportions. Only cells with non-missing categorical assignments were included in the corresponding kappa computations.

#### Spectral clustering analysis

Spectral clustering was performed using custom MATLAB scripts adapted from previously published methods (*52*). 2-sec segments of Ca^2+^ traces aligned to the onset of grasp (±1 sec time window) were extracted from individual cells and averaged across reaches. These mean segments were concatenated across control and MitoPark mice, and across cell types and age in a single matrix. Each segment was then z-scored using MATLAB’s *zscore*() function to normalize activity across cells so that clustering was driven by the temporal dynamics of activity patterns (i.e., temporal shape of the response) rather than absolute response magnitude. Dimensionality reduction was performed using principal component analysis (PCA), and the first 6 principal components (PCs) were retained, based on the elbow of the explained variance curve. A k-nearest neighbor affinity matrix was then constructed in PCA space and used to compute the graph Laplacian for spectral embedding. Clustering was then performed using k-means, with the optimal number of clusters determined by the silhouette score. Cluster stability was assessed by subsampling and quantified using the adjusted Rand index. Cluster identities were finally mapped back to experimental group, cell subtype, and age for subsequent analysis. The proportion of cells in each cluster was also calculated, and Chi-square tests were used for group comparisons.

#### Binomial logistic regression

Using custom MATLAB scripts, for each mouse, the denoised Ca^2+^ traces were z-scored using MATLAB’s *zscore()* function as described above. The traces were then aligned to timestamped grasp onset as previously described, and segments of 2 seconds around grasp onset were extracted as reach segments (see peri-event analysis for details). For comparison, randomly sampled non-overlapping segments of neural activity of identical duration (2 seconds) were drawn from periods without reaching, matched in number to reach segments (non-reach segments). Mice with recordings of fewer than 5 reaches at any age were excluded from this analysis. The extracted reach and non-reach segments were then concatenated across neurons and time to form reach-locked feature vectors. Reach versus non-reach segments were decoded using binomial logistic regression with 5-fold cross-validation. Within each training fold, dimensionality reduction was performed using PCA, retaining components that explained at least 85% of the variance. The classifier was fit on the training data using a logit link with Jeffreys-prior regularization, then applied to the held-out test data. Model performance was evaluated on held-out data using balanced accuracy and confusion matrices. Regression coefficients were back-projected from PCA space to the original neuron–time feature space and averaged across neurons within each cluster for each mouse for each week. This resulted in a time series for each mouse for each cluster for each week. Then separate generalized additive mixed models (GAMM) with restricted maximum likelihood for automatic smoothing were fit for each cluster within each week with a main effect of group, a smooth function of time, and a group-specific smooth interaction for MitoPark mice. Models additionally included a random intercept for mouse and autoregressive order-1 residual errors nested within mouse. This structure accounts for the correlation between observations from the same mouse that are close together in time. We then calculated the point-wise 95% confidence bands for the fitted smooth curves.

### Immunofluorescent staining and confocal imaging

At the end of recordings, mice were anesthetized with Ketamine (100 mg/kg) + Xylazine (20 mg/kg) and transcardially perfused with 1X phosphate-buffered saline (PBS) for 5 min followed by 4% paraformaldehyde (PFA) for 5 min. The mouse head with the implant was then placed into 4% PFA at 4°C for 24 hours. After careful removal of the implants, the brain was extracted from the skull, then placed into 4% PFA overnight at 4°C. Brains were then rinsed three times using PBS and sectioned (70 μm) using a VT1000s vibratome (Leica Biosystems, Deer Park, IL; RRID:SCR_016495). Slices containing M1 and the striatum were collected. Brain sections containing the striatum were used for tyrosine hydroxylase (TH) immunohistofluorescence studies. Brain sections were rinsed three times using phosphate-buffered saline (PBS), and incubated with 2% normal donkey serum in 0.5% Triton-X-100 PBS solution for 1 hour, followed by incubation with primary antibody against TH [Mouse, 1:2000, cat#: MAB318, MilliporeSigma; RRID: AB_2201528)] overnight at room temperature. The brain sections were rinsed three times with PBS and incubated with the secondary antibody Alexa Fluor 647 donkey anti-mouse IgG (1:500, cat#715-605-150, Jackson ImmunoResearch Labs, RRID: AB_2340862) for two hours at room temperature. After three rinses with PBS, brain sections were mounted on slides (H-1200, Vector Laboratories) and coverslipped. Details of the immunofluorescent staining protocol can be found on Protocols.io: 10.17504/protocols.io.b5s5q6g6. Immunofluorescence images were collected using a Nikon A1R Confocal Laser Scanning Microscope (RRID:SCR_020317). TH expression in the striatum and M1 was imaged with a 20x objective lens and quantified in ImageJ (NIH, https://imagej.net/, RRID: SCR_003070). TH immunofluorescence was quantified in the striatum and the cortex, and the striatal TH immunoreactivity was expressed as the ratio of striatum/cortex (Fig. S1). This was done in 3 separate sections for each mouse and averaged. Details of confocal imaging and TH quantification can be found on Protocols.io: 10.17504/protocols.io.3byl4jmxzlo5/v1 and 10.17504/protocols.io.n2bvj85qngk5/v1.

To register the location of lens implants in brain atlas coordinates, brain slices with implantation tracks were imaged using either a slide scanner (ZEISS Axio Scan.Z1 Slide Scanner, RRID:SCR_020927) or a confocal microscope (Leica SP8 lightning confocal microscope, RRID:SCR_018169). Images were then processed using the EBRAINS service for image registration (QuickNII and VisuAlign, https://ebrains.eu/service/quicknii-and-visualign, RRID:SCR_022818). Real AP, ML and DV coordinates of the prism lens implant were thus identified and compared to target implantation coordinates.

### Statistics

All data analyses were performed using the Inscopix IDEAS platform, GraphPad Prism 10 (Version 10, GraphPad Software, http://www.graphpad.com, RRID: SCR_002798), and custom scripts in MATLAB (MathWorks) and R (R Core Team). R packages lme4, emmeans, qgam, and mgcv were used. Figures were created using Adobe Illustrator (version 2024, https://www.adobe.com/products/illustrator.html, RRID:SCR_010279). Detailed statistics for each experiment can be found in the corresponding sections above, figure legends, and the raw data tables. Data were reported as mean and standard error of the mean (SEM). Statistically significant differences were identified at * p < 0.05, ** p < 0.01, *** p < 0.001, **** p < 0.0001.

## Acknowledgments

We thank the ASAP Team Wichmann for constructive discussion throughout this project. We also thank Gabriel Simms and Jessica Lugo for their participation in the preprocessing of calcium and behavioral videos.

## Author Contributions

Conceptualization: H.D.C., T.W., H.Y.C.

Methodology: H.D.C., D.B., B.R., H.Y.C.

Investigation: H.D.C., D.B., S.S., L.C., Z.W., J.J., B.R., H.Y.C.

Visualization: H.D.C., Z.W., J.J., B.R., H.Y.C.

Supervision: T.W., H.Y.C.

Writing—original draft: H.D.C., H.Y.C.

Writing—review & editing: H.D.C., B.R., T.W., H.Y.C.

## Competing interests

All authors declare they have no competing interests.

### Data, code, and materials availability

The raw and analyzed data generated throughout this study, as well as the Matlab and R codes used for data analysis, have been deposited and are publicly available from: 10.5281/zenodo.23734033 and 10.48324/dandi.002042/0.261028.2212

## Supplementary Materials

Figs.S1-10

Tables S1-2

