## Supplementary figures for "Emergence of Task-Related Motor Cortical Dysfunction in Mice with Progressive Parkinsonism": Supplementary figures and figure legends v5.pdf

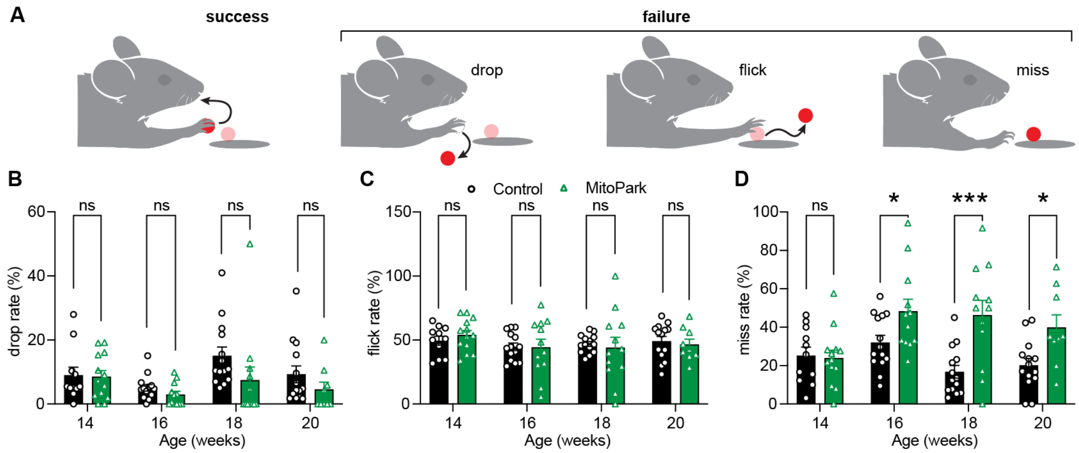


**Supplementary Figure 1. Behavioral changes across different categories of failed reaches. A)** Illustrative definition of different categories of reaches. **B-D)** Rate of failed reaches, specifically drop (B), flick (C) and miss (D) reaches, in MitoPark mice and littermate controls. Linear mixed-effects model. Each data point represents one mouse. MitoPark = 13, 12, 13, 12 mice and control = 11, 14, 14, 14 mice at 14, 16, 18 and 20 weeks, respectively. Detailed numbers and statistics are available in the Source Data Table for all figures.

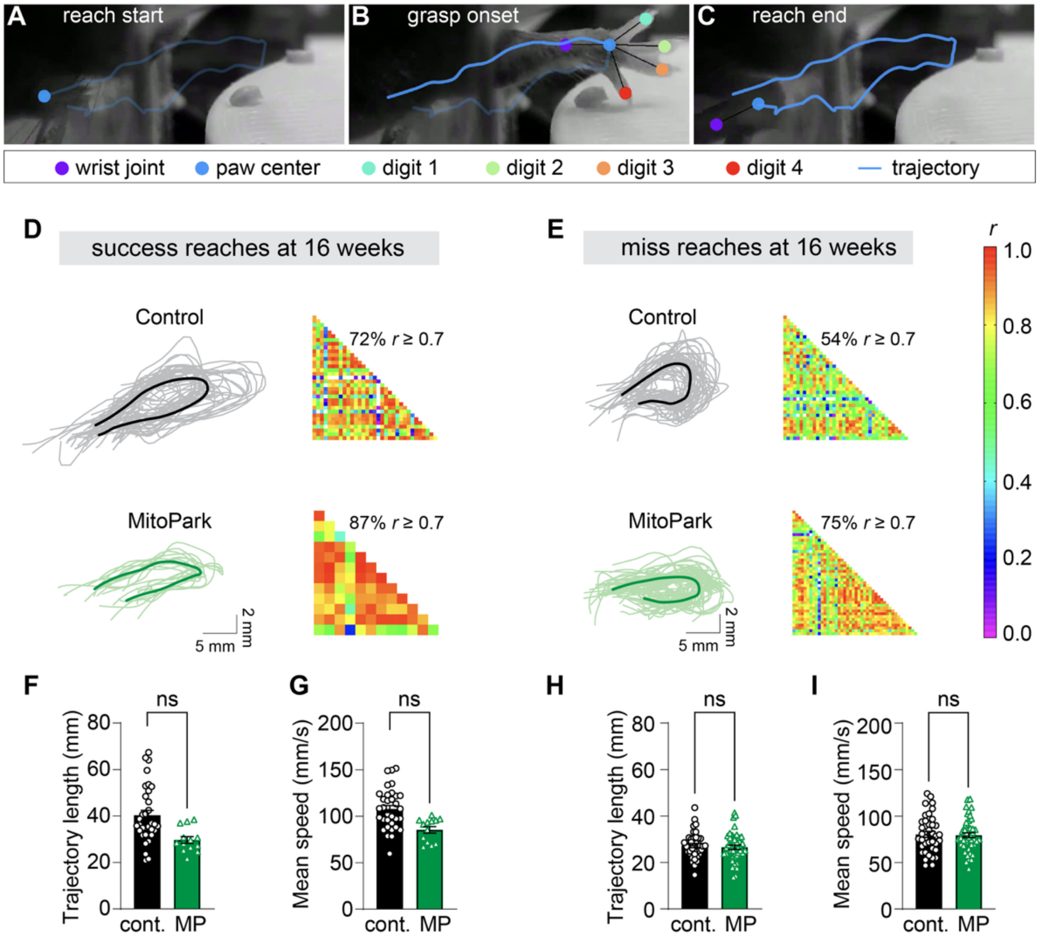


**Supplementary Figure 2. Paw movement kinematics of control and MitoPark mice. A-C)** Example trajectory of a success reach, showing the start of the movement at the metal perch (A), the moment of grasp onset (B), and the end of the reach back to the starting position (C). B shows how the paw was labeled and the skeleton built on DeepLabCut, the wrist and digits serving as anchors for the paw center tracking. **D-E)** Correlation analysis of successful reaches (D) and missed reaches (E) at 16 weeks. Each panel shows the resampled trajectories of the reaches (left), the correlation matrix (right) with the percentage of highly correlated pairs of reaches (Pearson's $r \geq 0.7$). White squares in the matrices represent negative values. Successful reaches: Control n/N = 33 reaches/3 and MitoPark n/N = 13/3 reaches/mice. Missed reaches: Control n/N = 39 reaches /3 mice and MitoPark n/N = 45 reaches/3 mice. **F-G)** The length of reach trajectory and mean speed of paw movement during success reaches. **H-I)** The same as (F-G) but for missed reaches. Each data point in (F-I) represents one reach. Control: n/N = 140, 282, 181 reaches from 3 mice, and MitoPark: n/N = 67, 208, 47 reaches from 3 mice at 14, 16 and 18 weeks, respectively.

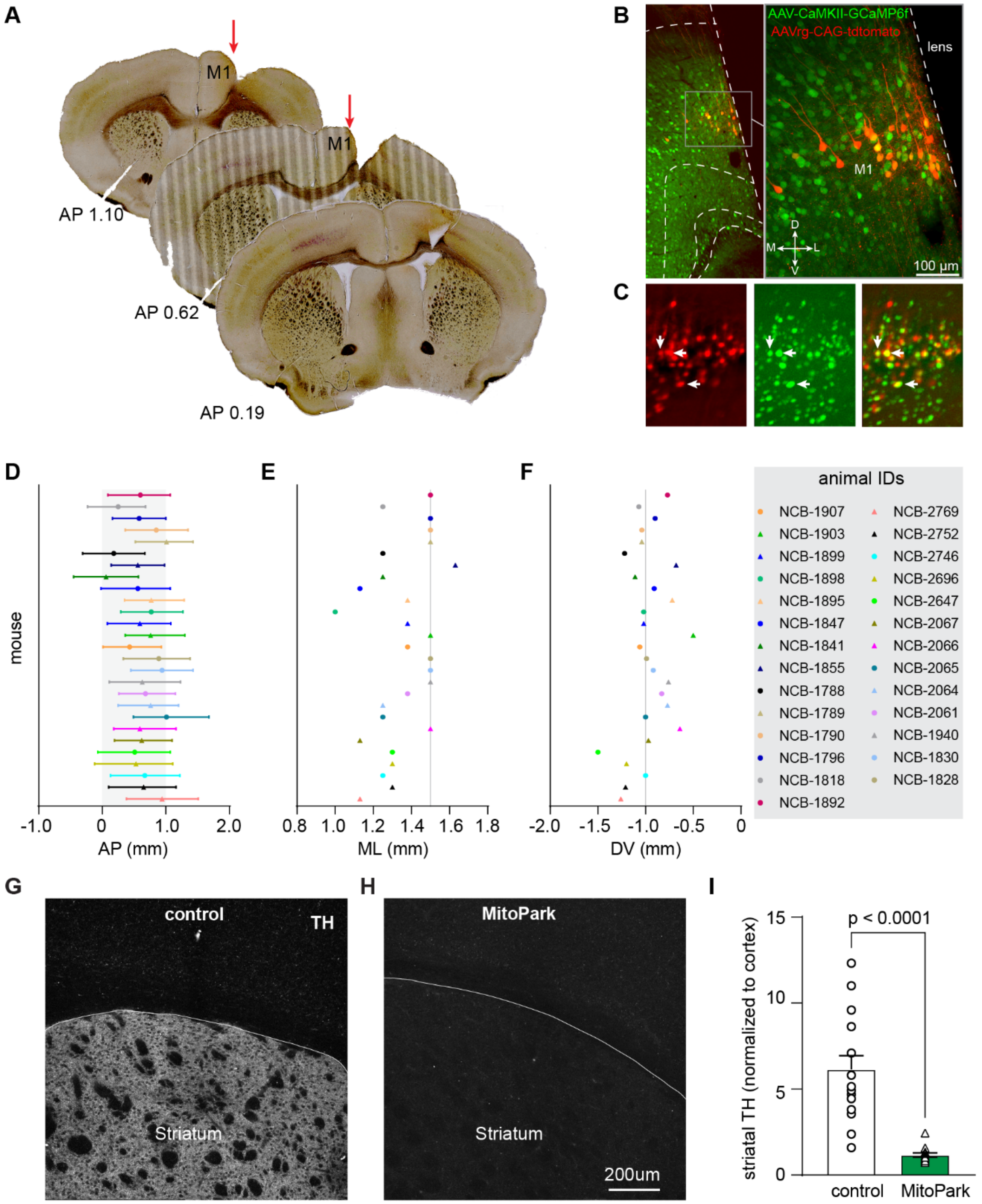


**Supplementary Figure 3. Histological validation of implantation site and degeneration of the nigrostriatal dopaminergic projection in MitoPark mice. A)** Representative images used for atlas registration. **B-C)** Representative *post hoc* confocal images (B) and *in vivo* field of view (FoV) of miniScope showing the laminar location of the center of FoV. Arrows in (C) highlight co-labeled CSp neurons in M1. **D-F)** The antero-posterior (D), medio-lateral (E), and dorso-ventral

(F) coordinates of the GRIN lens implant of all mice used in this study. The gray shaded area in D and the gray vertical lines in E and F represent the target coordinates. Individual animals are listed on the right. **G-H)** Representative images of striatal TH staining in control and MitoPark mice. **I)** Striatal TH quantification. Each data point represents one mouse. Control = 6.14 ± 0.8, N = 15 mice, MitoPark = 1.17 ± 0.13, N = 13 mice, $p < 0.0001$. Mann-Whitney U test.

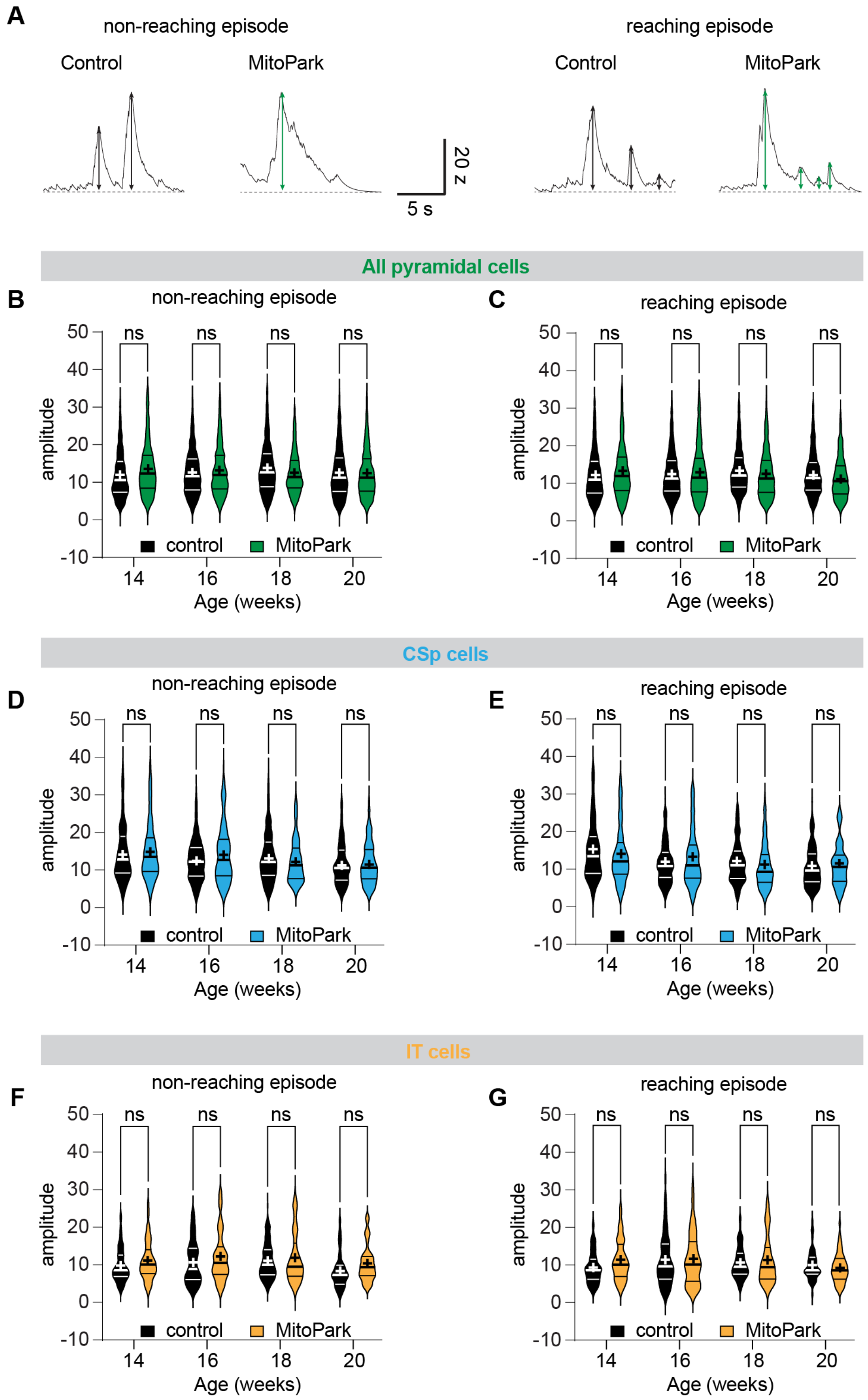


**Supplementary Figure 4. Amplitude of $Ca^{2+}$ events of M1 pyramidal neurons. A)** Representative $Ca^{2+}$ transients from a control mouse (gray) and a MitoPark mouse (green), during non-reaching epochs (left) and reaching epochs (right). **B-G)** Amplitude $Ca^{2+}$ events averaged over non-reaching (B, D, F) and reaching epochs (C, E, G) of M1 pyramidal cells (B-C), CSp cells (D-E) and IT cells (F-G). Horizontal lines indicate median and IQR, cross (+) symbols indicate the mean. Violin plots represent cells. Linear mixed-effects model. (B) Control n/N (cells/mice)= 1203/11, 1385/13, 1699/14 and 1610/14, MitoPark n/N (cells/mice) = 1074/13, 1459/12, 2072/11 and 1677/12, at 14, 16, 18 and 20 weeks, respectively. (C) Control n/N (cells/mice) = 1191/11, 1354/13, 1712/14 and 1612/14, MitoPark n/N (cells/mice) = 1052/13, 1431/12, 1298/11 and 1268 /9 at 14, 16, 18 and 20 weeks, respectively. (D) Control n/N (cells/mice) = 83/5, 189/6, 242/6 and 177/6, MitoPark n/N (cells/mice) = 138/7, 175/7, 171/7 and 191/7, at 14, 16, 18 and 20 weeks of age, respectively. (E) Control n/N (cells/mice) = 76/5, 179/6, 240/6 and 177/6, MitoPark n/N (cells/mice) = 129/7, 166/7, 134/6 and 87/4, at 14, 16, 18 and 20 weeks, respectively. (F) Control n/N (cells/mice) = 78/6, 63/7, 131/7 and 50/7, MitoPark n/N (cells/mice) = 88/6, 81 /5, 108/6 and 58/5, at 14, 16, 18 and 20 weeks, respectively. (G) Control n/N (cells/mice) = 78/6, 63/7, 132/8 and 49/8, MitoPark n/N (cells/mice) = 89/6, 82 /5, 51/3 and 58/5, at 14, 16, 18 and 20 weeks, respectively. Detailed numbers and statistics are available in the Source Data Table.

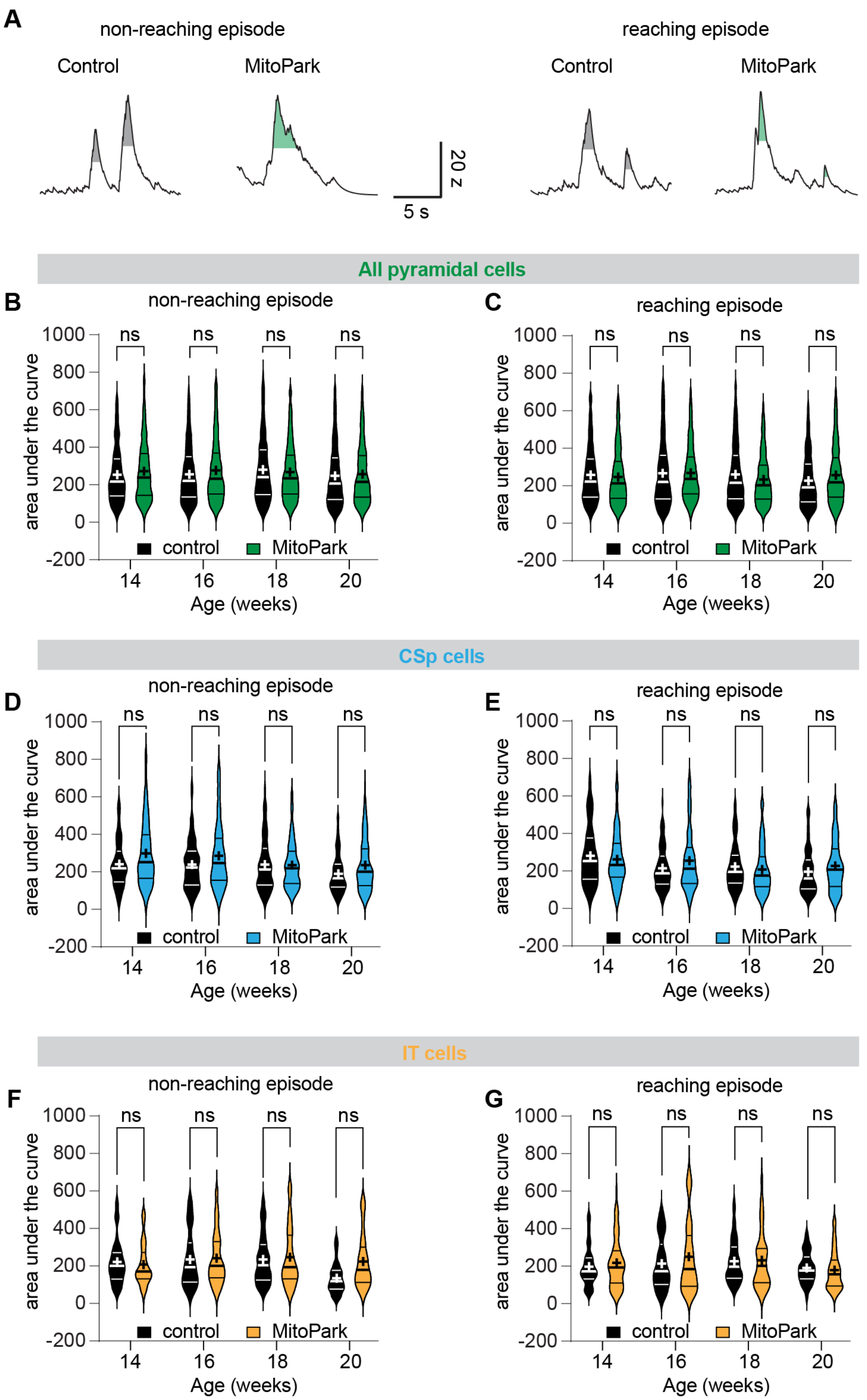


**Supplementary Figure 5. Area under the curve of $Ca^{2+}$ traces of M1 pyramidal neurons. A)** Representative $Ca^{2+}$ transients from a control mouse (gray) and a MitoPark mouse (green), during non-reaching epochs (left) and reaching epochs (right). **B-G)** Area under the curve of $Ca^{2+}$ events averaged over non-reaching (B, D, F) and reaching epochs (C, E, G) of M1 pyramidal cells (B-C), CSp cells (D-E) and IT cells (F-G). Horizontal lines indicate median and IQR, cross (+) symbols indicate the mean. Violin plots represent cells. Linear mixed-effects model. (B) Control n/N = 1203 cells/11 mice, 1385 cells/13 mice, 1699 cells/14 mice and 1610 cells/14 mice, MitoPark n/N = 1074 cells/13 mice, 1459 cells/12 mice, 2072 cells/11 mice and 1677 cells/12 mice, at 14, 16, 18 and 20 weeks, respectively. (C) Control n/N = 1191 cells/11 mice, 1354 cells/13 mice, 1712 cells/14 mice and 1612 cells/14 mice, MitoPark n/N = 1052 cells/13 mice, 1431 cells/12 mice, 1298 cells/11 mice and 1268 cells/9 mice at 14, 16, 18 and 20 weeks, respectively. (D) Control n/N = 83 cells/5 mice, 189 cells/6 mice, 242 cells/6 mice and 177 cells/6 mice, MitoPark n/N = 138 cells/7 mice, 175 cells/7 mice, 171 cells/7 mice and 191 cells/7 mice, at 14, 16, 18 and 20 weeks of age, respectively. (E) Control n/N = 76 cells/5 mice, 179 cells/6 mice, 240 cells/6 mice and 177 cells/6 mice, MitoPark n/N = 129 cells/7 mice, 166 cells/7 mice, 134 cells/6 mice and 87 cells/4 mice, at 14, 16, 18 and 20 weeks, respectively. (F) Control n/N = 78 cells/6 mice, 63 cells/7 mice, 131 cells/7 mice and 50 cells/7 mice, MitoPark n/N = 88 cells/6 mice, 81 cells/5 mice, 108 cells/6 mice and 58 cells/5 mice, at 14, 16, 18 and 20 weeks, respectively. (G) Control n/N = 78 cells/6 mice, 63 cells/7 mice, 132 cells/8 mice and 49 cells/8 mice, MitoPark n/N = 89 cells/6 mice, 82 cells/5 mice, 51 cells/3 mice and 58 cells/5 mice, at 14, 16, 18 and 20 weeks, respectively. Detailed numbers and statistics are available in the Source Data Table.

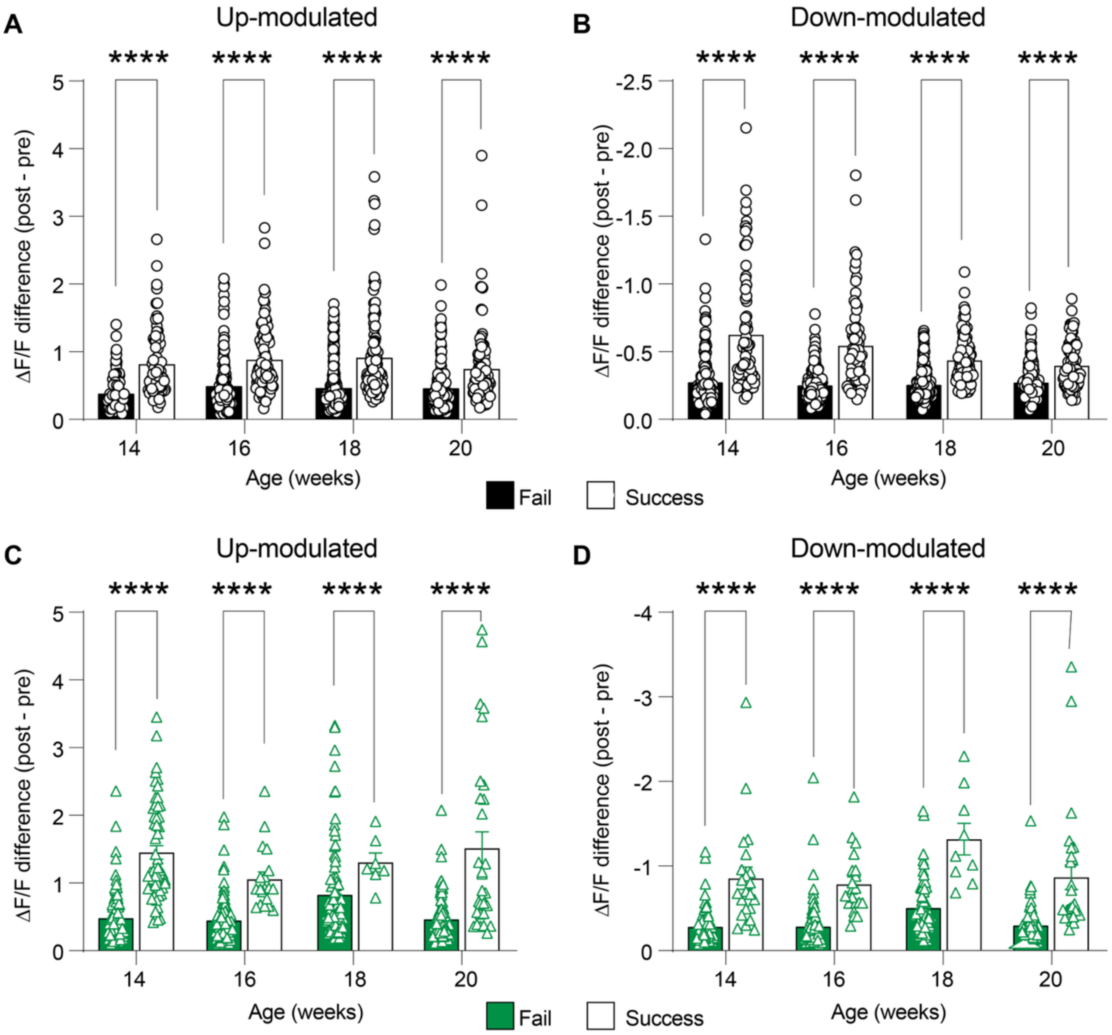


**Supplementary Figure 6. Reach outcome-specific modulation of M1 pyramidal neurons. A-D)** Summary results of the magnitude of modulation for up- (A, C) and down-modulated (B, D) pyramidal cells when their activity is aligned to failed vs successful reaches, within control (A-B) and MitoPark (C-D) mice. Each data point represents one cell. Linear mixed-effects model. (A) Upmodulated, control: fail, n/N = 155/11, 171/13, 234/13, 186/14 cells/mice and success, n/N = 77/11, 90/12, 109/13, 97/10 at 14, 16, 18 and 20 weeks, respectively. (B) Downmodulated, control: fail, n/N = 198/11, 215/12, 313/13, 230/13 and success, n/N = 73/9, 69/12, 100/12, 85/10 cells/mice at 14, 16, 18 and 20 weeks, respectively. (C) Upmodulated, MitoPark: fail, n/N = 102/11, 129/11, 96/10, 99/8 and success, n/N = 54/11, 19/4, 7/2, 30/6 cells/mice. (D) Downmodulated, MitoPark: fail, n/N = 97/12, 120/11, 65/10, 81/9 cells/mice, and success, n/N = 22/8, 20/5, 9/2, 27/5 cells/mice at 14, 16, 18 and 20 weeks, respectively.

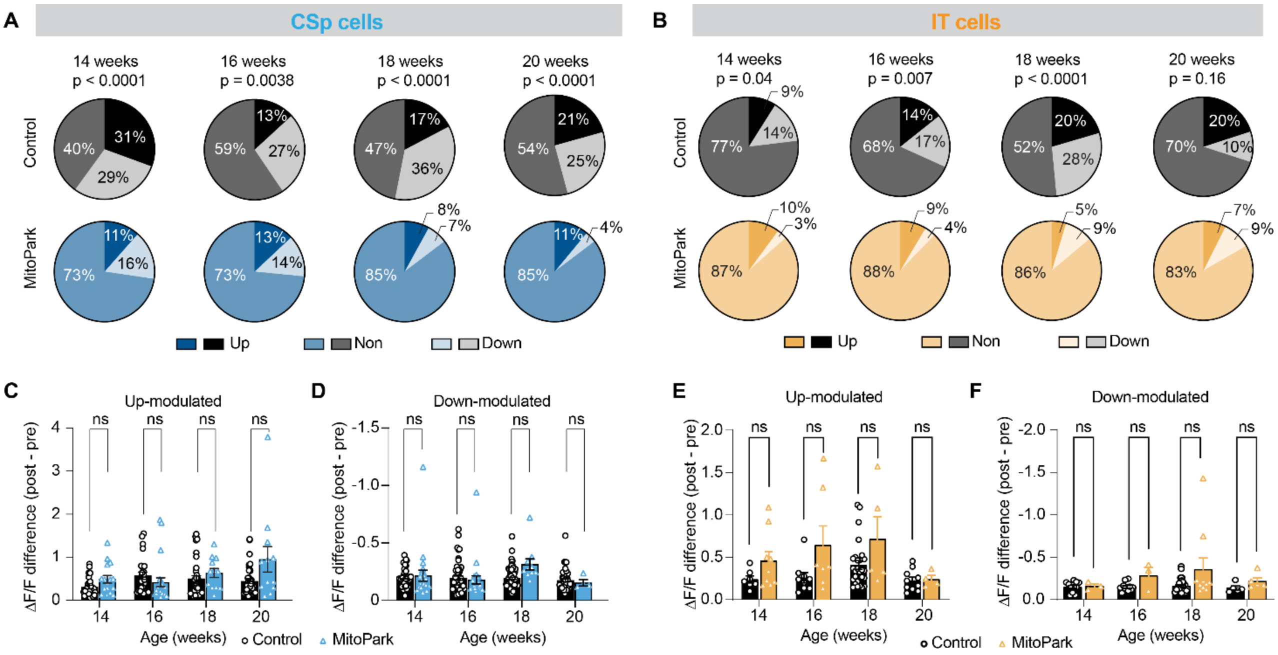


**Supplementary Figure 7. Movement modulation of CSp and IT neurons in MitoPark mice and controls. A-B)** Proportions of different categories of CSp (A) and IT (B) neurons at each time point. Chi-square test to compare differences. **C-F)** Summary results of the magnitude of modulation for up- (C, E) and down-modulated (D, F) cells within CSp (C, D) and IT (E, F) neurons. Each data point represents one cell. Linear mixed-effects model, followed by multiple comparisons with Benjamini-Hochberg correction. (C) Upmodulated CSp: Control n/N = 26/5, 25/5, 42/6, 37/6 cells/mice and MitoPark n/N = 16/6, 23/7, 12/5, 12/4 cells/mice at 14, 16, 18 and 20 weeks, respectively. (D) Downmodulated CSp: Control n/N = 25/5, 52/6, 87/6, 45/6 cells/mice and MitoPark n/N = 22/5, 242/6, 10/5, 4/1 cells/mice at 14, 16, 18 and 20 weeks, respectively. (E) Upmodulated IT: Control n/N = 7/5, 9/5, 27/5, 10/6 cells/mice and MitoPark n/N = 9/3, 7/3, 5/3, 4/3 cells/mice at 14, 16, 18 and 20 weeks, respectively. (F) Downmodulated IT: Control n/N = 11/3, 11/5, 37/7, 5/2 cells/mice and MitoPark n/N = 3/2, 3/2, 10/3, 5/3 cells/mice at 14, 16, 18 and 20 weeks, respectively. Detailed numbers and statistics are available in the Source Data Table.

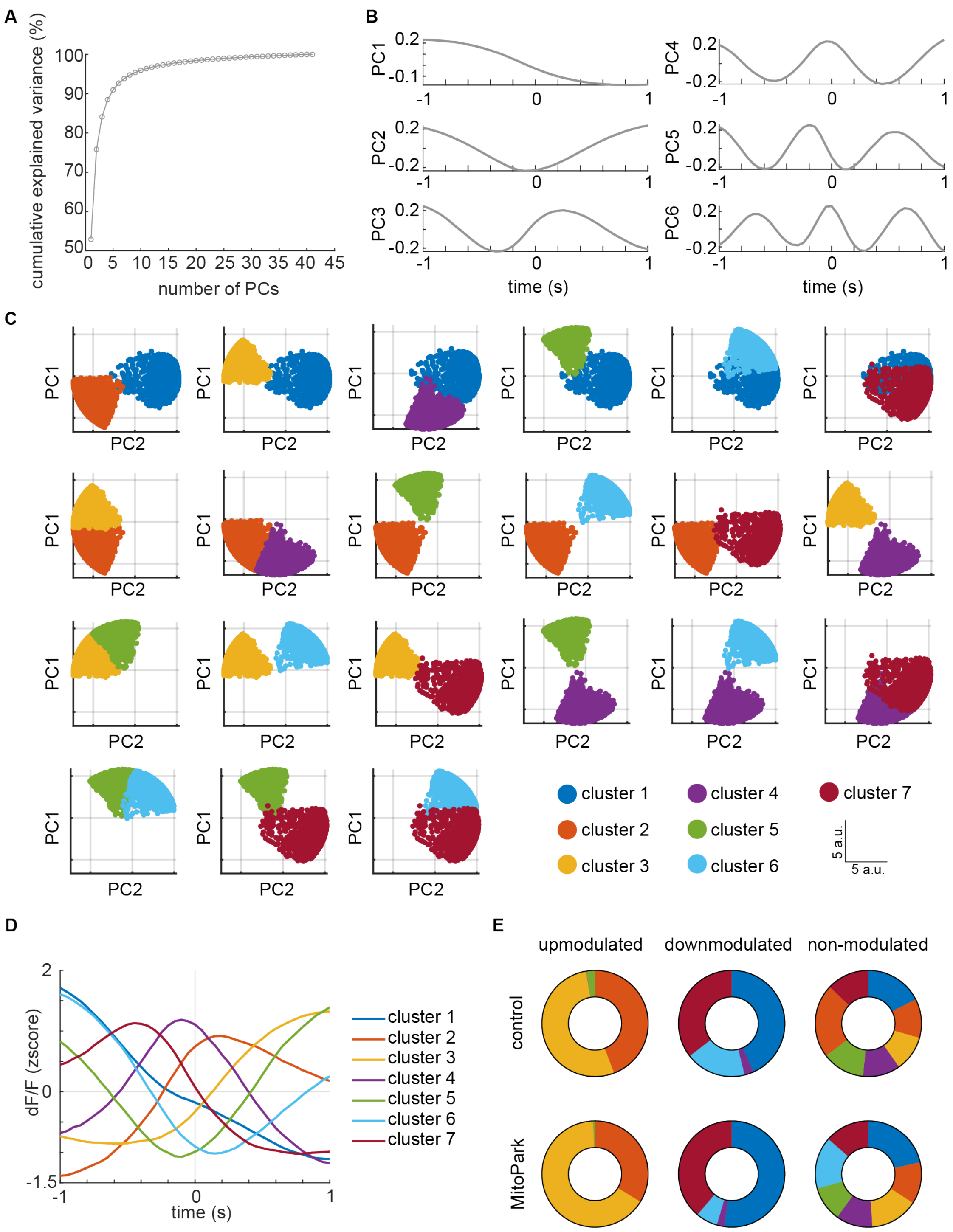


**Supplementary Figure 8. Spectral clustering analysis of M1 pyramidal neurons. A)** Cumulative explained variance plot. Elbow method was used, and 6 PCs were retained. **B)** Plots of the weights of the first 6 PCs. **C)** Cells projected onto 2D PC space to show clusters separation. Each subplot shows PC1 versus PC2 for one pair of clusters. **D)** Mean traces of each cluster showing the different functional temporal profiles identified, averaged across age and experimental groups for representative purposes. **E)** Overlap between neurons identified as upmodulated (left), downmodulated (middle), and non-modulated (right) and cluster identity in control (top) and MitoPark mice (bottom), pooled across ages. Colors denote cluster identity and are consistent across panels.

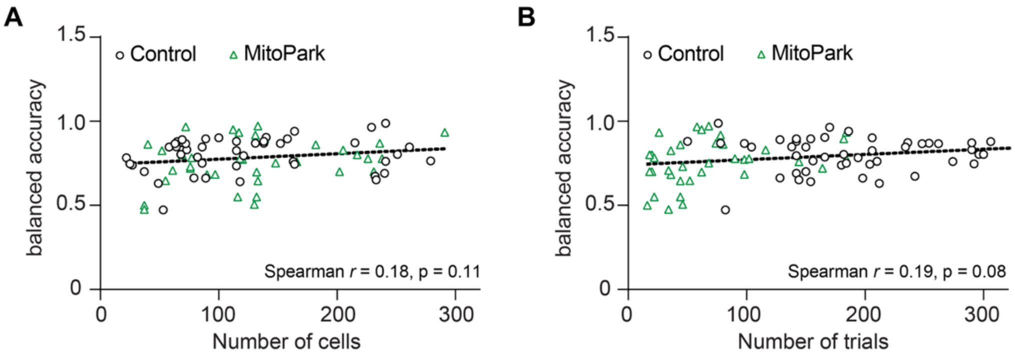


**Supplementary Figure 9.** Correlation between number of cells (A) and number of trials (B) with balanced accuracy obtained by the binomial logistic regression, showing no correlation between the variables.

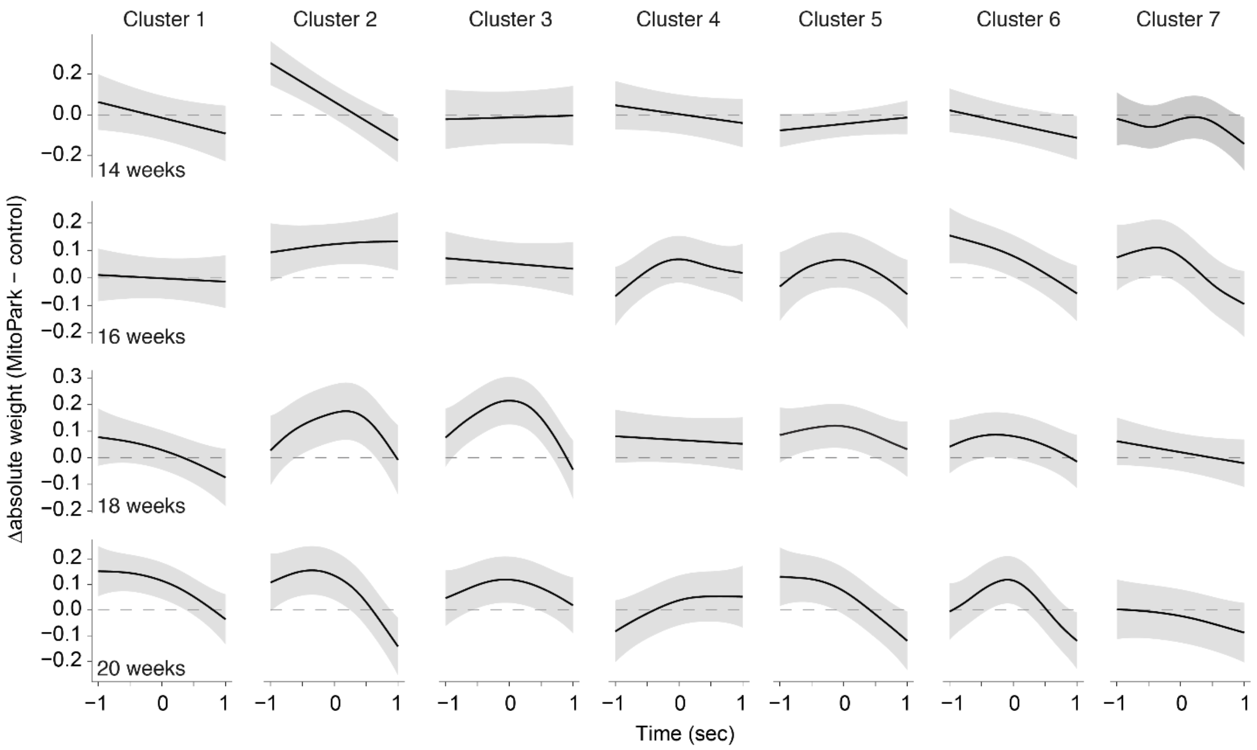


**Supplementary Figure 10. Difference between the reach encoding curves in control and MitoPark mice.** For each week and each cluster, a GAMM with autocorrelated errors was fit to predict the absolute temporal weights from time, with separate smooths fit for MitoPark and control. The difference curves were obtained by subtracting the smooth for control from MitoPark. Gray bands indicate point-wise 95% confidence bands.
